# Loss of Ku’s DNA end binding activity affects telomere length via destabilizing telomere-bound Est1 rather than altering TLC1 homeostasis

**DOI:** 10.1101/600197

**Authors:** Laramie D. Lemon, Danna K. Morris, Alison A. Bertuch

## Abstract

*Saccharomyces cerevisiae* telomerase, which maintains telomere length, is comprised of an RNA component, TLC1, the reverse transcriptase, Est2, and regulatory subunits, including Est1. The Yku70/Yku80 (Ku) heterodimer, a DNA end binding (DEB) protein, also contributes to telomere length maintenance. Ku binds TLC1 and telomere ends in a mutually exclusive fashion, and is required to maintain levels and nuclear localization of TLC1. Ku also interacts with Sir4, which localizes to telomeres. Here we sought to determine the role of Ku’s DEB activity in telomere length maintenance by utilizing *yku70-R456E* mutant strains, in which Ku has reduced DEB and telomere association but proficiency in TLC1 and Sir4 binding, and TLC1 nuclear retention. Telomere lengths in a *yku70-R456E* strain were nearly as short as those in *ykuΔ* strains and shorter than in strains lacking either Sir4, Ku:Sir4 interaction, or Ku:TLC1 interaction. TLC1 levels were decreased in the *yku70-R456E* mutant, yet overexpression of TLC1 failed to restore telomere length. Reduced DEB activity did not impact Est1’s ability to associate with telomerase but did result in decreased association of Est1 with the telomere. These findings suggest Ku’s DEB activity maintains telomere length homeostasis by preserving Est1’s interaction at the telomere rather than altering TLC1 levels.

## INTRODUCTION

Telomeres are nucleoprotein complexes that cap the natural ends of linear eukaryotic chromosomes^1^. In capping, telomeres protect chromosome ends from degradation and recombination, thus preserving genomic integrity. As a result of the end-replication problem, terminal telomeric DNA is loss with each round of semi-conservative DNA replication^2^. Across eukarya, with the notable exception of dipterans, telomeric DNA consists of tandem repeats of simple G-rich sequence^3, 4^. The length of telomeric DNA, including sequence lost due to the end-replication problem, can be maintained by the enzyme telomerase, a specialized reverse transcriptase that has an integral RNA component that provides the template for *de novo* synthesis of telomere repeats onto native chromosome ends^5^.

The unicellular organism *Saccharomyces cerevisiae* has proven to be a powerful model system to study telomerase and telomere length regulation^6, 7^. In *S. cerevisiae*, telomerase minimally consists of an RNA component, TLC1,^8^ and a catalytic component, Est2, both of which are required for telomerase activity *in vitro* and *in vivo*^9, 10^. Additional subunits of telomerase include Est1, which binds a stem-loop structure of TLC1^11^, and Est3, which associates with the telomerase complex via binding sites on Est1 and Est2^12, 13^. Est1 is involved in the recruitment of telomerase to the telomere during late S phase through its association with Cdc13, a single-stranded DNA binding protein that binds the terminal 3’ overhang of the telomeric G-rich strand with high affinity^14–19^. In addition, there is evidence to support a second role for Est1 subsequent to telomerase recruitment^19, 20^, which is often referred to as activation^14^. The molecular nature of telomerase activation, however, remains undetermined. The role of Est3 in telomere maintenance similarly remains unclear, but it has been hypothesized that part of Est1's activation function involves the recruitment of Est3 to the telomere^21^. Unlike TLC1 and Est2, Est1 and Est3 are not required when telomerase activity is assayed in cell lysates^9, 22^. However, like TLC1 and Est2, Est1 and Est3 are required for telomerase action at native chromosome ends. Loss of any of these four major components of telomerase results in progressive telomere shortening and cellular senescence^8, 23, 24^.

Additional factors have been linked to telomere length maintenance in *S. cerevisiae*. One such factor is the evolutionarily conserved Ku heterodimer, which consists of Ku70 and Ku80 subunits (known as Yku70 and Yku80 in *S. cerevisiae*). Ku binds DNA ends with high affinity and independently of sequence using a ring-like structure^25^. Not surprisingly, Ku plays a major role in DNA double strand break repair, specifically in the non-homologous end joining (NHEJ) pathway, where, upon induction of a DNA double strand break, Ku rapidly binds the DNA ends, providing protection and a platform for the recruitment of NHEJ factors^26^.

Remarkably, Ku also associates with telomeres^27^, which are normally immune to NHEJ, and is involved in several aspects of telomere structure and function. In this work, we focus on Ku's role in telomere length maintenance, which is reflected by yeast strains lacking either of the Ku subunits manifesting very short telomeres^28, 29^. In contrast to strains lacking telomerase components, Ku deficient strains have stably short telomeres and do not senesce.

Several models have been put forth to explain the impact of Ku on telomere length maintenance. One model involves Ku’s impact on the subcellular localization and steady state levels of TLC1. Ku associates with a specific stem-loop structure on TLC1^30, 31^; this binding is required for the nuclear retention of TLC1 as cells lacking Ku or Ku:TLC1 interaction (such as *yku80-135i* or *tlc1∆-48* mutants) fail to retain TLC1 in the nucleus^32, 33^. Additionally, steady state levels of TLC1 are decreased in *yku∆*, *yku80-135i*, and *tlc1∆-48* mutants, although the mechanism responsible for this reduction remains unknown^34, 35^. Regardless of mislocalization and reduction in levels, a sufficient amount of TLC1 must be retained in the nucleus in Ku-deficient cells to circumvent continued shortening and prevent senescence.

An additional mechanism by which Ku has been proposed to promote telomere length maintenance involves its interaction with the silent information regulator Sir4. Sir4 localizes to telomeres via its association with 15-20 Rap1 molecules that are bound directly along the ~300 basepairs (bp) of telomeric DNA^36, 37^. Sir4 also binds Ku and this interaction is an important pathway for telomerase recruitment to the telomere^14, 38^. In support of this, Est2 enrichment at telomere VI-R is greatly reduced in strains bearing mutations in key Ku80 binding residues on Sir4^14^. While telomere length is only slightly shortened in these or *sir4∆* mutants, an important role for Ku:Sir4 interaction in telomere maintenance was demonstrated by the finding that strains bearing certain mutations that impair both Ku:Sir4 and Est1:Cdc13 interactions are senescent whereas those with the single mutations are not^14^. Interestingly, Ku separation-of-function mutants that no longer bind Sir4 and have impaired telomerase recruitment have telomere lengths that are only slightly shorter than wild type, whereas *yku∆* strains have drastically short telomeres^14, 39, 40^.

A third model for the role of Ku in telomere length maintenance is via its effects on Est1. Ku promotes the association of both Est1 and Est2 with the telomere^41^. Although Est1 and Est2 associate with the telomere in a mutually dependent fashion^42^, Ku impacts the association of Est1 to the telomere even when Est2’s recruitment to the telomere is mediated by its fusion with Cdc13^33^. Thus, it is unlikely that the reduction of Est1’s telomere association in a Ku null strain is due to reduced Est2 at the telomere. Ku is also found in a complex with Est1, an association that is dependent on the presence of TLC1. Additionally, the requirement for Ku in telomere length maintenance is bypassed when Est1 is tethered to the telomere via Cdc13 but not when Est2 is tethered^33^. Thus, Ku’s role in telomere length maintenance also likely involves the Est1 protein.

Notably, Ku’s engagement of the telomere end via its DNA end binding (DEB) activity is not employed in any of these models. Indeed, studies using purified Ku, linear DNA, and the TLC1 stem-loop to which Ku binds illustrated that Ku associates with these nucleic acids in a mutually exclusive fashion^43^. This was further supported by structural studies that demonstrated that the Ku:TLC1 stem-loop occupies a position on the heterodimer that would preclude Ku’s loading onto a DNA end when bound to TLC1^14, 25^. Nonetheless, Ku’s DEB activity is required for its influence on telomere length^44^. The *yku70-R456E* mutant, which has reduced binding to DNA and the telomere end, has stably short telomeres, despite binding TLC1 and retaining its localization in the nucleus like wild type strains^33, 44^. Additionally, the *yku70-R456E* mutant is proficient for Sir4 binding suggesting that loss of Ku’s DEB activity alone is sufficient to cause the short telomere phenotype^44^.

In this study, we aimed to further characterize the role of Ku’s DEB activity in telomere length homeostasis using the *yku70-R456E* mutant of Ku. We found shorter telomeres in a *yku70-R456E* strain than in strains bearing mutations disrupting the telomerase-Ku-Sir4 pathway. We also found decreased TLC1 levels in the *yku70-R456E* mutant. However, TLC1 overexpression failed to restore telomere length, despite increasing TLC1 levels in the nucleus, suggesting another consequence of impaired Ku DEB results in telomere shortening. Consistent with this, we found that, while Est1’s ability to bind TLC1 was not impacted in the *yku70-R456E* mutant, its association with the telomere was greatly impaired. Additionally, the interaction of Est1:Yku80 was reduced, whereas soluble Est1:Cdc13 complexes were increased. These data lead us to propose that, by loading onto the telomere end, Ku contributes to telomere length maintenance by preserving Est1’s interaction with the telomere.

## RESULTS

### Telomeres in the *yku70-R456E* mutant are shorter than other Ku mutants and comparable to *yku∆* strains

We first sought to directly compare the telomere lengths of a mutant yeast strain that had impaired interaction between Ku and telomere ends (*yku70-R456E*)^44^, with those with impaired interaction between Sir4 and Yku80 (*sir4∆, yku80-L111R*, and *yku80-L115A*)^14^, impaired interaction between TLC1 and Yku80 (*tlc1∆-48* and *yku80-135i*)^30, 31^, or lacked Ku altogether and, therefore, were deficient in all of these functions (*yku70∆* and *yku80∆*) (Fig. 1 and Table 1). We found the telomeres in the *yku70-R456E* mutant were much shorter than in the *sir4∆*, *yku80-L111R,* and *yku80-L115A* strains. Because the Yku70-R456E/Yku80 heterodimer can still bind Sir4^44^, the shorter telomere lengths in the *yku70-R456E* mutant strain indicates that Ku’s role in telomere length maintenance is not entirely mediated through the Ku:Sir4 interaction. In addition, we found the telomeres in the *yku70-R456E* mutant were also shorter than in *tlc1∆-48* (as previously reported^44^) and *yku80-135i* strains. While the shorter telomere lengths in *yku80-135i* and *tlc1∆-48* mutants as compared to *sir4∆, yku80-L111R*, and *yku80-L115A* mutants has been explained by the additional loss of TLC1 nuclear retention^14^, this could not explain the even shorter telomeres in the *yku70-R456E* mutant, as TLC1 is still retained the nucleus in this mutant strain^33^.

**Table 1.** Telomere length in Ku null and separation-of-function mutants.

|  | Mutant | Telomere length<br>relative to WT (bp) |
| --- | --- | --- |
| Null | *yku70*Δ | -199 ± 7 |
| Null | *yku80*Δ | -190 ± 10 |
| Defective for DEB | *yku70-R456E* | -161 ± 10 |
| Defective for Ku:TLC1 interaction | *tlc1*Δ48 | -110 ± 14 |
| | *yku80-135i* | -108 ± 33 |
| Defective for Ku:Sir4 interaction | *sir4*Δ | -45 ± 7 |
| | *yku80-L111R* | -30 ± 13 |
| | *yku80-L115A* | -28 ± 14 |

Values based on n=3

**Figure 1.**
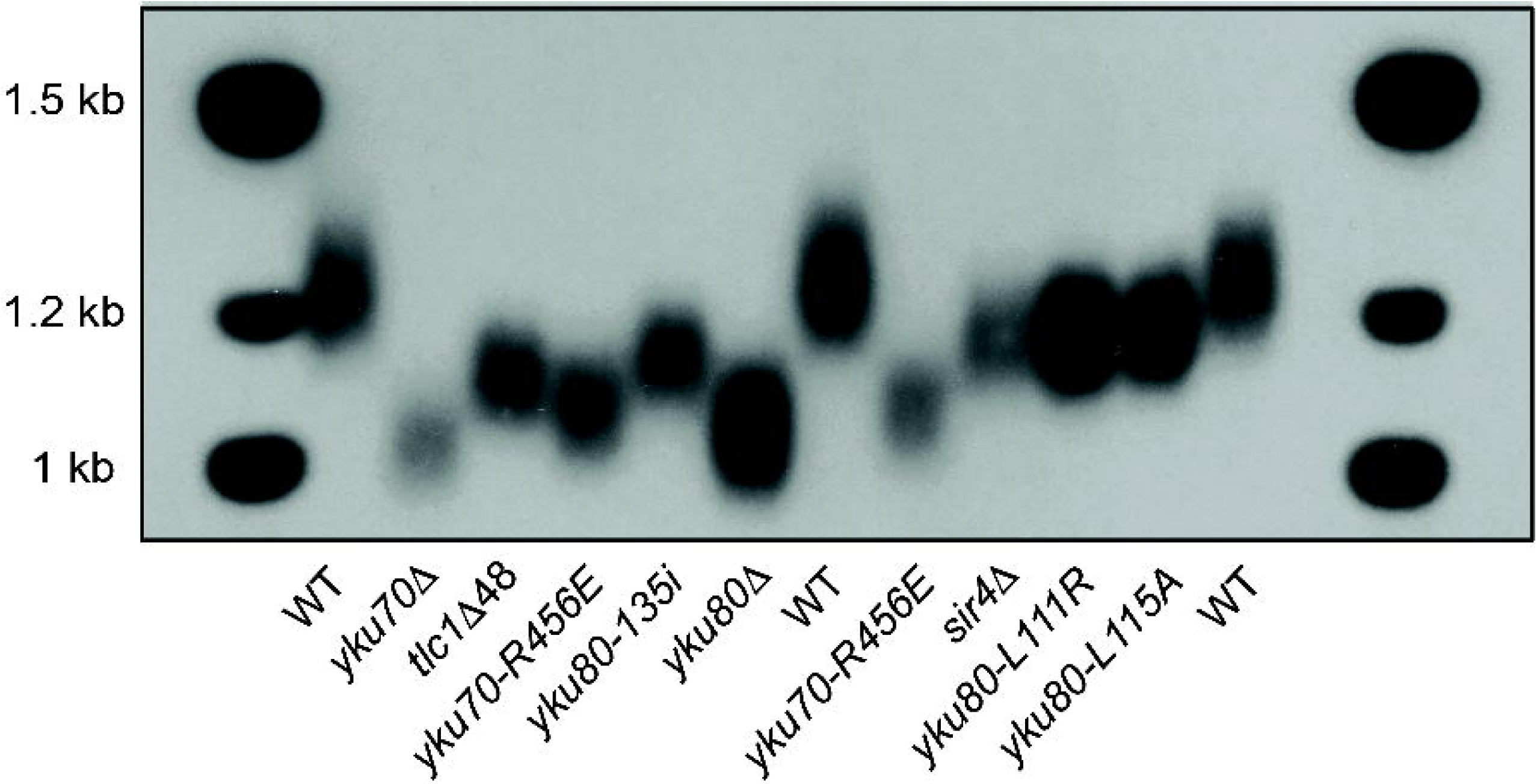
Telomere lengths of the *yku70-R456E* mutant compared to other mutants. Telomere length analysis by Southern blotting of strains as indicated. The full-length blot is presented in Supplementary Figure S6.

### The short telomere phenotype of the *yku70-R456E* mutant is not due to reduced TLC1 levels

TLC1 levels are decreased in strains lacking Ku or Ku:TLC1 interaction although the mechanism underlying this reduction remains unknown^34, 35^. This led us to ask if a decrease in steady state TLC1 levels was the cause of the short telomere phenotype in the Ku DEB mutant. We first determined the relative amount of TLC1 in asynchronous *yku70-R456E* cells as compared to wild type cells using reverse transcriptase quantitative polymerase chain reaction (RT-qPCR) and northern blotting. We found TLC1 levels were reduced in the *yku70-R456E* mutant by approximately 60%, similar to the reduction displayed in *yku∆* strains (Fig. 2). Furthermore, when we expressed *YKU70* in a *yku70∆* strain, TLC1 levels increased nearly two-fold whereas expression of the *yku70-R456E* mutant had no impact compared to the empty vector control (Supplementary Fig. S1). Because this mutant of Ku with reduced DEB also has less TLC1, this suggests that Ku’s DEB activity is required for TLC1 stability. Moreover, because TLC1 nuclear retention and binding are preserved in the yku70-R456E (unlike the *yku80-135i* or *yku80∆* mutants), this indicates that nuclear retention by Ku is not sufficient to maintain TLC1 levels.

**Figure 2.**
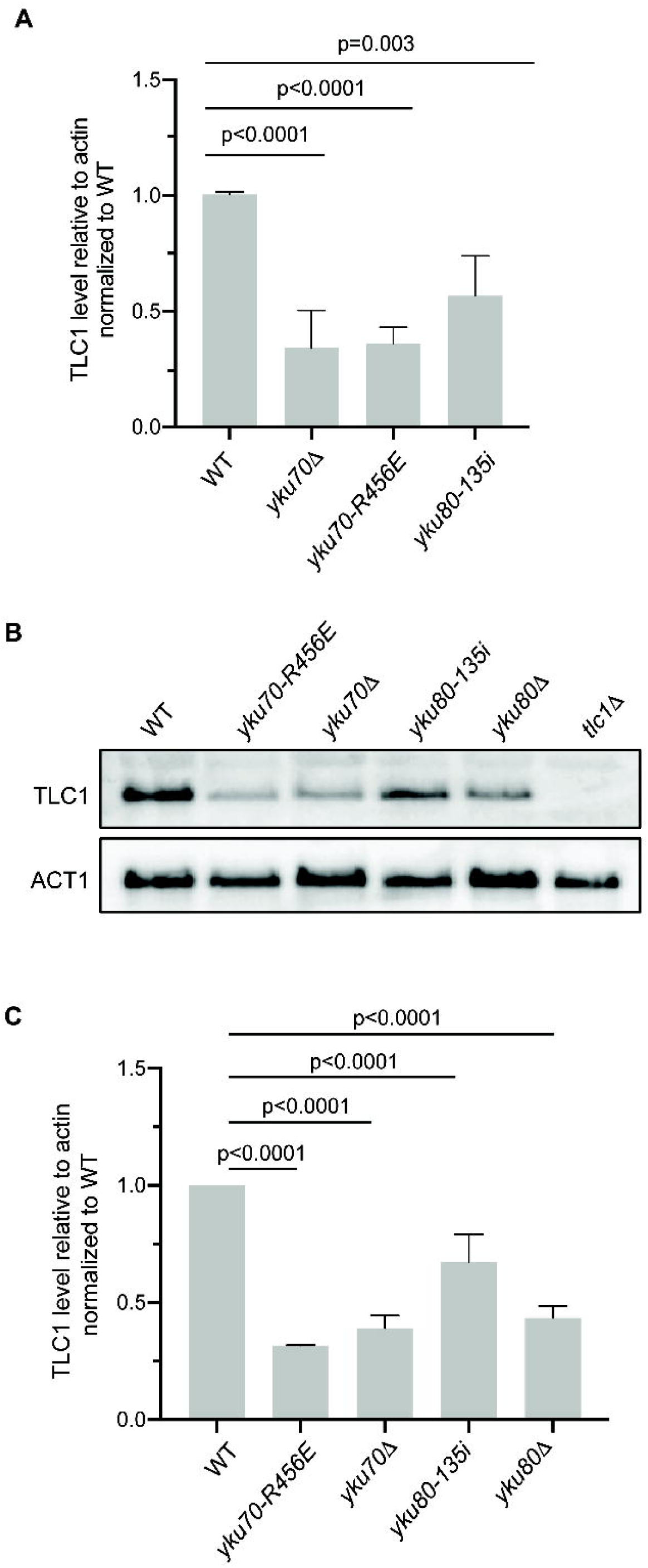
TLC1 levels are decreased in Ku mutants. (A) Quantification of TLC1 RNA in asynchronous wild type (WT), *yku70-R456E*, *yku70*Δ, and *yku80-135i* strains by RT-qPCR in four independent experiments. TLC1 levels were quantified relative to actin RNA and normalized to WT. Error bars represent ± 1 standard deviation (SD). p values adjusted for multiple comparisons are shown. (B) Analysis of TLC1 levels by northern blotting of asynchronous WT, *yku70-R456E*, *yku70*Δ, *yku80-135i*, *yku80*Δ and *tlc1*Δ strains. Blots were also probed with ACT1 for loading. Full-length blots are presented in Supplementary Figure S7. (C) Quantification of three independent northern blotting experiments. Error bars represent ± 1 (SD). p values adjusted for multiple comparisons.

Having identified the decrease in TLC1, we considered the possibility that this could contribute to the shortened telomeres in the *yku70-R456E* mutant. Because Ku’s ability to interact with TLC1 was retained in the *yku70-R456E* mutant^44^, we reasoned that overexpression of TLC1 might restore telomere length in these cells. Therefore, we overexpressed TLC1 by expression from the *ADH1* promoter on a 2 micron plasmid, which is maintained at high copy number, and assessed telomere length over successive streakouts using Southern blotting. Despite marked TLC1 overexpression (Figs. 3A,B), telomeres remained short in the *yku70-R456E* mutant (Fig. 3C). Importantly, the lack of rescue of telomere length was not due to a failure of overexpressed TLC1 to localize to the nucleus (Fig. 3D). Thus, the short telomere phenotype that results from a decrease in the DEB activity of Ku is not due reduced TLC1 levels. These findings suggest that Ku’s DEB activity itself is inherently important for telomere length maintenance.

**Figure 3.**
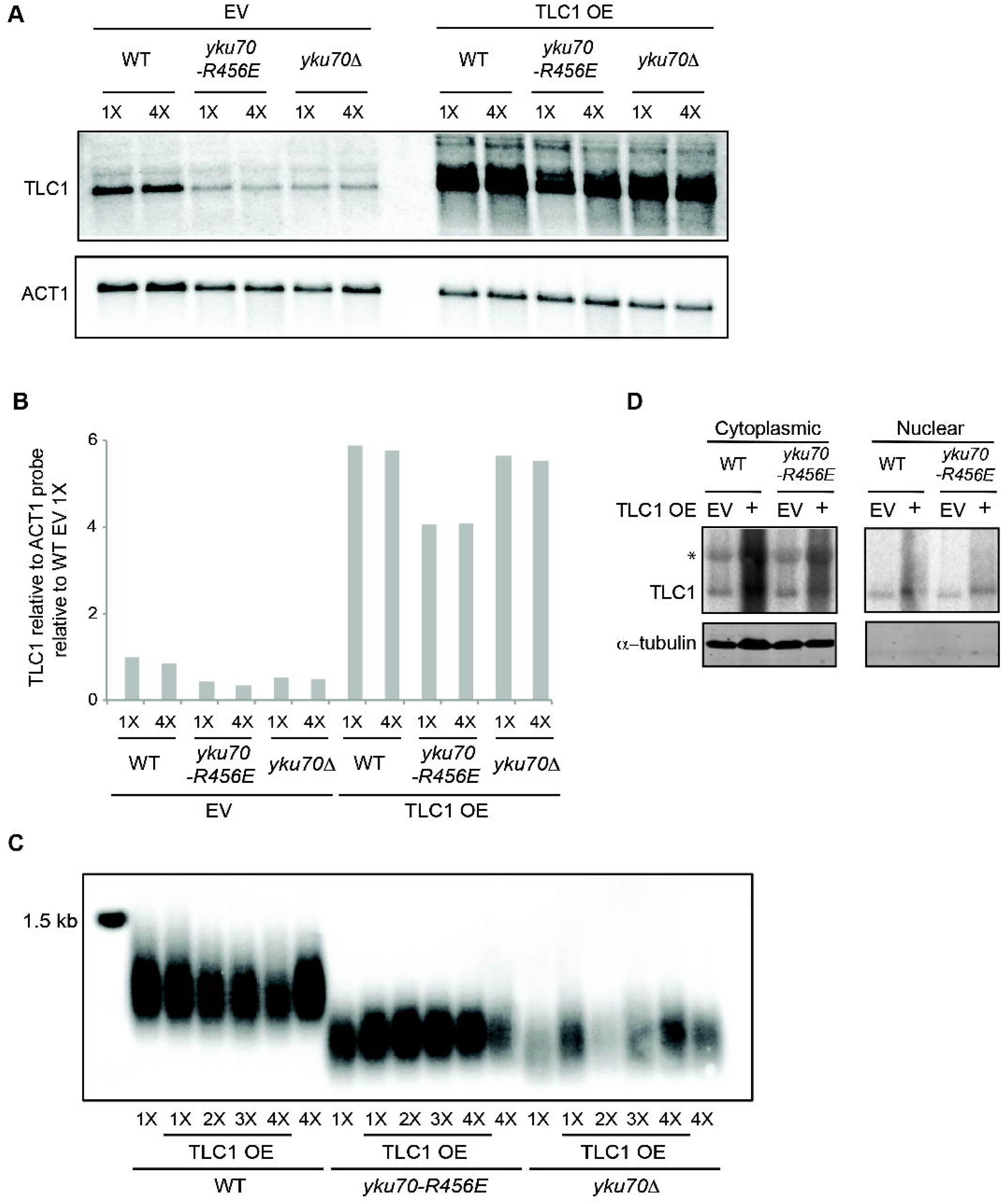
The short telomere phenotype of the *yku70-R456E* mutant cannot be rescued by overexpressing TLC1. (A) TLC1 RNA overexpression (OE) was confirmed by northern blotting of WT, *yku70-R456E*, and *yku70*Δ strains transformed with a 2 micron *TLC1* plasmid (TLC1 OE) or a 2 micron empty vector (EV) following 1X and 4X streak outs. Blots were also probed with ACT1 for loading. Full-length blots are presented in Supplementary Figure S8. (B) Quantification of northern blot shown in A. TLC1 levels were quantified relative to ACT1 RNA and normalized to WT EV 1X streakout cells. (C) Telomere length analysis by Southern blotting of 1X-4X serial streakouts of WT, *yku70-R456E*, and *yku70*Δ strains harboring 2 micron TLC1 OE or EV plasmids. The full-length blot is presented in Supplementary Figure S9. (D) Northern blot probed for TLC1 RNA in cytoplasmic and nuclear fractions (top) and western blot analyzed for tubulin, which is a cytoplasmic protein. The absence of tubulin the nuclear fraction indicates the nuclear prep was free of cytoplasmic contamination. The full-length northern blot and western blot are presented in Supplementary Figure S10. *presumed nonspecific band (see retention of a similarly migrating band in the *tlc1Δ* lane of full-length northern blot of Fig. 2B which is presented in Supplementary Fig. S7).

In support of this notion, we previously found that telomeres are extensively and progressively elongated in *yku70-R456E* and *yku70∆* mutants when Est1, but not Est2, is tethered to the telomere via fusion with Cdc13^33^. While a possible reason for this differential effect on telomere elongation may be that the Cdc13-Est1 fusion alone restores TLC1 levels in these mutants, we found TLC1 amounts were unchanged in *yku70-R456E* and *yku70∆* strains expressing the Cdc13-Est1 fusion when assessed by northern blot and RT-qPCR and over serial streakouts (Fig. 4 and Supplementary Fig. S2). These findings further suggest that it is effects on Est1 rather than TLC1 that underlie the telomere length maintenance defects in the *yku70-R456E* mutant.

**Figure 4.**
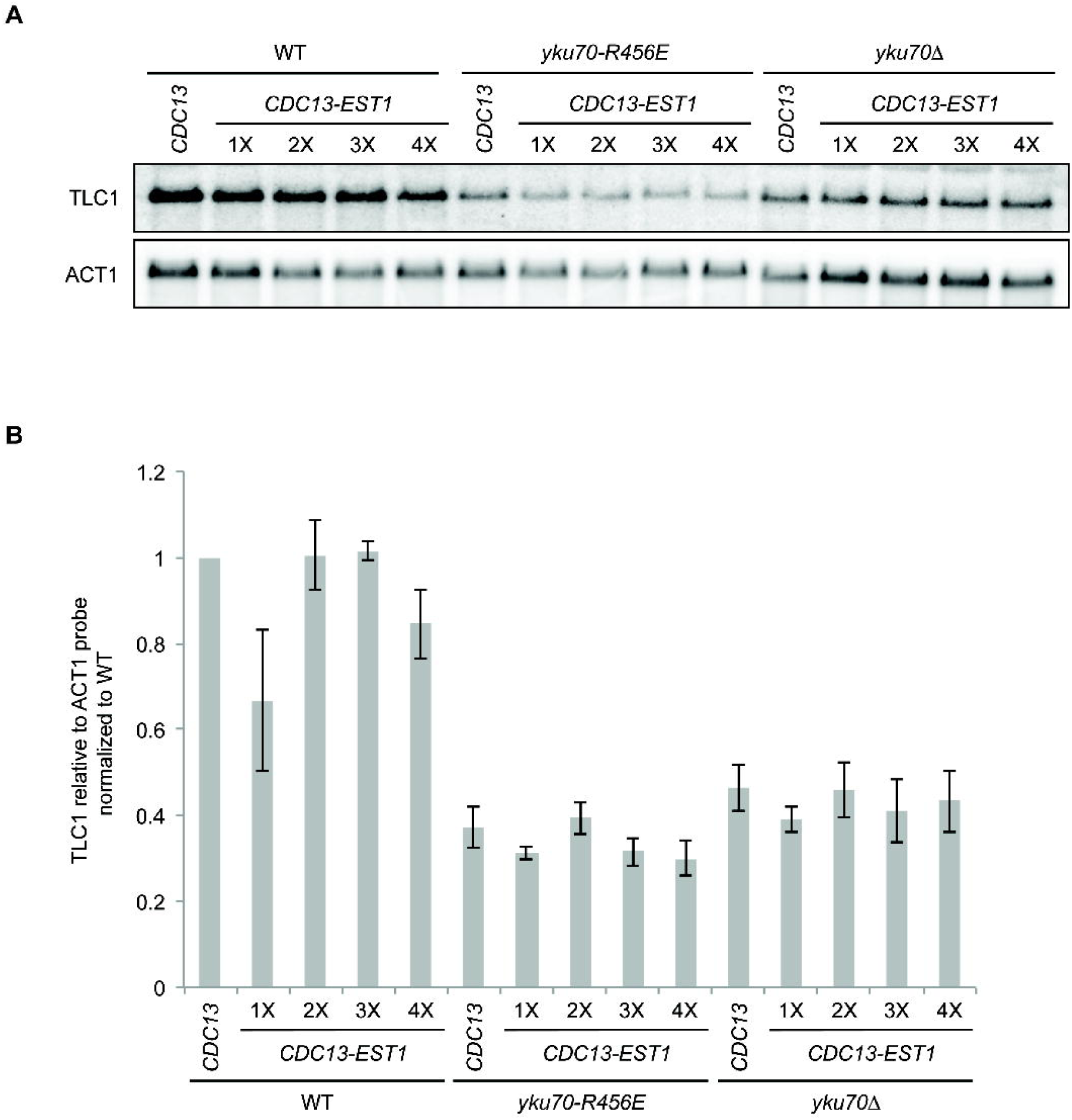
TLC1 levels are not stabilized in Ku mutant strains expressing *CDC13-EST1* fusions. (A) Analysis of TLC1 levels by northern blotting of 1X-4X serial streakouts of *cdc13*Δ (WT), *cdc13*Δ *yku70-R456E* (*yku70-R456E*), and *cdc13*Δ *yku70*Δ(*yku70*Δ) strains expressing either Cdc13 or a Cdc13-Est1 fusion. Blots were also probed with ACT1 for loading. Full-length blots are presented in Supplementary Figure S11. (B) Quantification of the mean from three independent northern blotting experiments. Error bars represent ± standard error of the mean (SEM).

### Ku’s DEB activity influences its association with Est1 and Est1’s association with telomere

We next turned our attention to the role of Ku’s DEB activity on Ku’s association with Est1 and on Est1’s association with TLC1 and the telomere. We previously demonstrated that Ku’s association with Est1, as assayed by co-immunoprecipitation, was dependent on Ku’s interaction with TLC1^33^. As anticipated given the reduction in TLC1 (Fig. 2), there was less Est1 in Yku80 immunoprecipitates in the *yku70-R456E* mutant strain (Fig. 5A,B). However, the association was not rescued by overexpression of TLC1 (Fig. 5C,D), despite increased TLC1 in the *yku70-456E* mutant’s nucleus when TLC1 was overexpressed (Fig. 3D) and retention of the Yku70-R456/Yku80 heterodimer’s ability to bind TLC1^44^.

**Figure 5.**
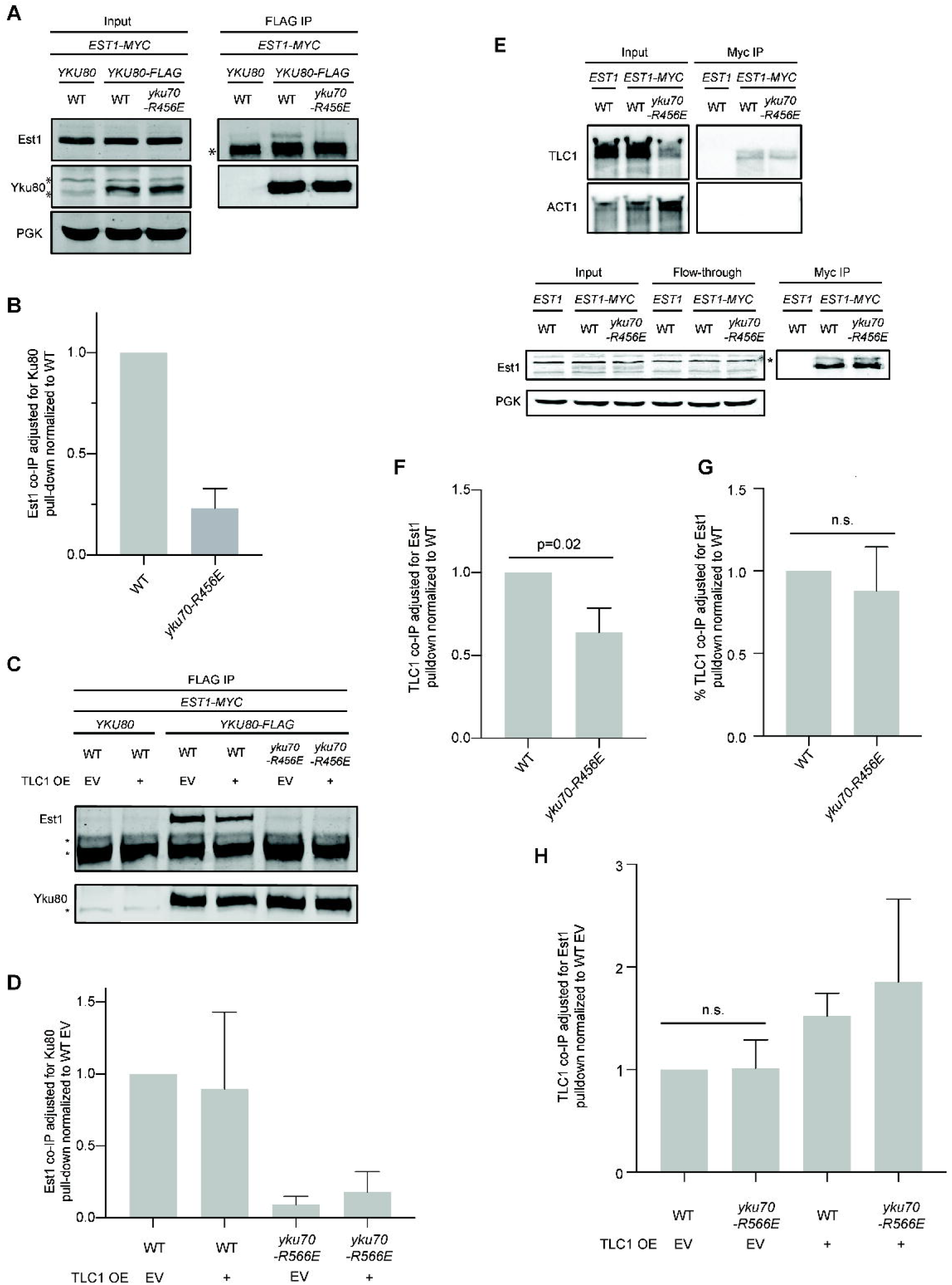
Ku’s DEB activity influences Est1:Ku but not Est1:TLC1 interaction. (A) Co-immunoprecipitation (co-IP) of Est1-myc with Yku80-FLAG. Anti-FLAG immunoprecipitations were performed with whole cell lysate of indicated strains. Immunoprecipitates (IP) and inputs were analyzed by western blotting with α-myc (Est1) and α-FLAG (Yku80). “ * ” indicates non-specific bands in western blots. Inputs were also probed with α-PGK as a loading control. (B) Quantification represents the mean Est1 co-IP relative to Yku80-FLAG pull-down normalized to WT (n=3). Error bars represent ± SD. (C) As in (A) except only IPs are shown with indicated strains transformed with either TLC1 OE or EV. (D) As in (B) except with normalization to WT EV. (E) Co-immunoprecipitation of TLC1 with Est1-myc. Anti-myc immunoprecipitations were performed with whole cell lysate of indicated strains. Isolated RNA was analyzed by northern blot and probed with a radiolabeled TLC1 specific probe (top). Blots were also probed with ACT1 for loading. Western blotting was performed to assess Est1-myc pull-down efficiency (bottom) “*” nonspecific band. (F) Quantification represents mean TLC1 RNA co-IP’d relative to Est1 pulldown normalized to WT based on three independent experiments. Error bars represent ± SD. (G) Quantification represents mean percent TLC1 RNA co-IP’d relative to Est1 pulldown normalized to WT based on the same three independent experiments as in (F). Error bars represent ± SD. n.s., nonsignificant. (H) Quantification of three independent experiments as in (E) except with indicated strains transformed with either TLC1 OE or EV. Quantification was performed as in (F) except p values were adjusted for multiple comparisons, all of which were n.s. Representative blots are shown in Supplementary Figure S3. Full-length blots are presented in Supplementary Figure S12 (A), S13 (C), S14 (E), and S15 (for Fig. S3 corresponding to H).

To determine whether Ku’s DEB activity impacted Est1’s interaction with TLC1, we examined TLC1 levels in Est1 immunoprecipitates in wild type and *yku70-R456E* mutant strains. We found that Est1:TLC1 interaction was reduced by approximately 40% in the *yku70-R456E* mutant compared to wild type (Fig. 5E,F). When normalized to the amount of TLC1 in the input, however, there was no difference in Est1:TLC1 interaction in the *yku70-R456E* mutant and wild type strains (Fig. 5G), suggesting the reduction in Est1:TLC1 interaction was due to a reduction in TLC1 levels in the Ku DEB mutant. As in the Est1:Yku80 interaction studies, we also examined the effect of TLC1 overexpression on Est1:TLC1 interaction in *yku70-R456E*. No difference in Est1:TLC1 interaction was observed in wild type versus *yku70-R456E* strains when transformed with EV or TLC1 OE (Fig. 5H and Supplementary Fig. S3). The reason for the different observations in the set of experiments without and with EV or TLC1 OE plasmids is unclear; however, taken together, they suggest Est1:TLC1 interaction is not impacted by Ku DEB.

Previous results indicated that the principal role for Ku in telomere length maintenance lies in its influence on Est1’s telomere association, in part because Ku promotes the association of Est1 to the telomere even when Est2 is tethered to the telomere via a Cdc13 fusion^33^. Therefore, we directly tested whether Ku’s DEB activity is required for Est1’s association with the telomere using telomere chromatin immunoprecipitation (ChIP). Like the *yku70∆* strain, we found a marked reduction of Est1 binding telomeric chromatin in the *yku70-R456E* mutant (Fig. 6). Taken together, these findings suggest that Ku’s DEB ability is important for maintaining telomere length homeostasis by preserving Est1’s interactions with the telomere.

**Figure 6.**
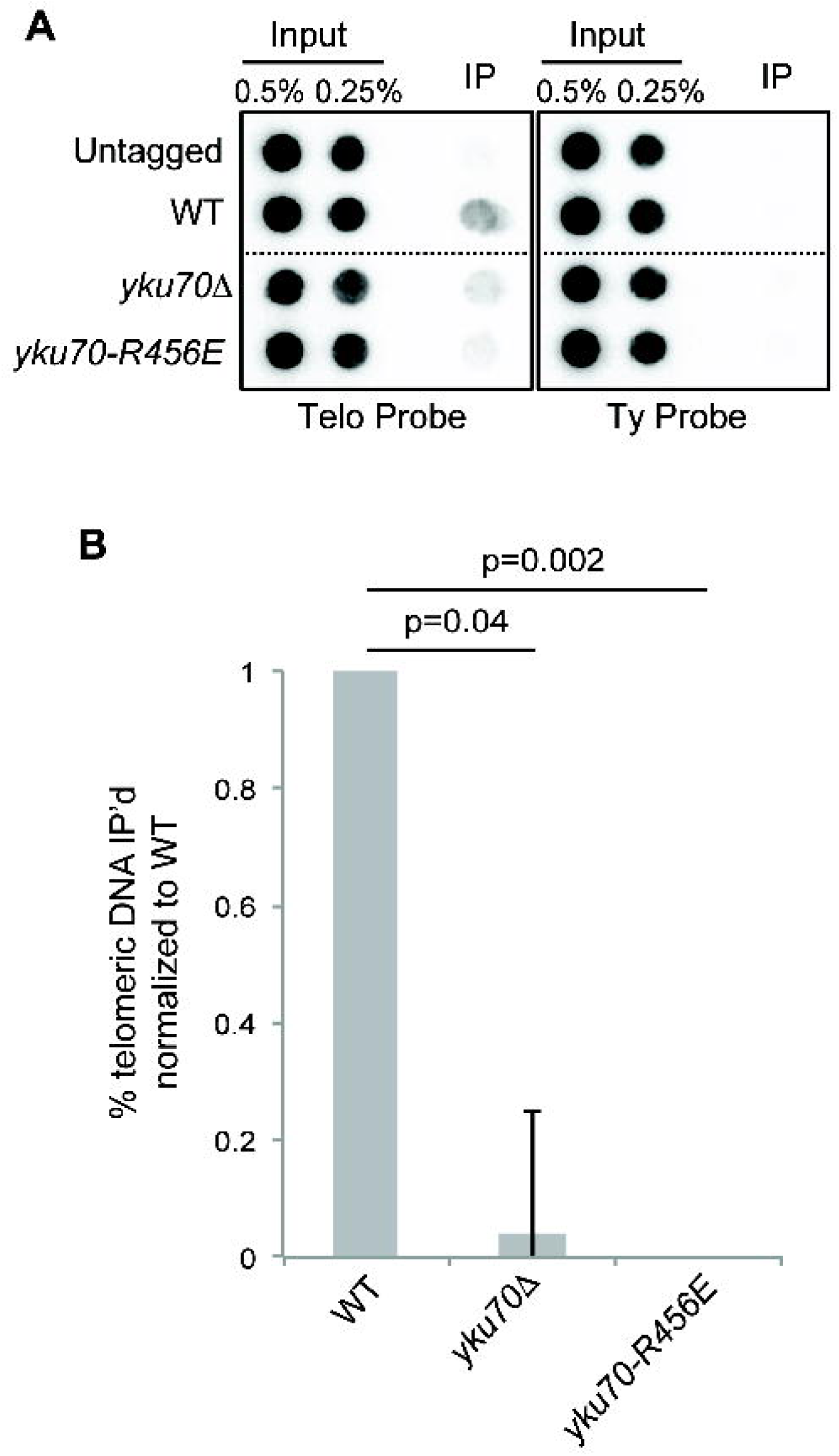
Ku DEB supports Est1’s telomere association. (A) ChIP of Est1-myc. Anti-myc immunoprecipitations were performed on formaldehyde crosslinked WT, *yku70∆*, or *yku70-R456E* strains. Isolated DNA was analyzed by dot blotting and probed with a radiolabeled telomere specific probe or TyB to assess nonspecific interaction. The lines indicate areas in which the image was cropped to remove nonrelevant samples. Full-length blots are presented in Supplementary Figure S16. (B) Quantification of the percent of telomeric DNA associated with Est1-myc normalized to WT for three independent experiments. Error bars represent ± SEM.

### Soluble Est1:Cdc13 complexes increase when Ku’s DEB is impaired

If Ku maintains telomere length by preserving Est1’s interactions with the telomere, we next reasoned that Ku might promote Est1’s binding to Cdc13. To test this, we performed co-immunoprecipitation of endogenously tagged Est1 and Cdc13 proteins in asynchronous strains. Interestingly, soluble Est1:Cdc13 complexes were increased in *yku70∆* and *yku80∆* strains compared to wild type (Fig. 7). This increase could be rescued by expressing *YKU80* from a centromeric plasmid (Supplementary Fig. S4).

**Figure 7.**
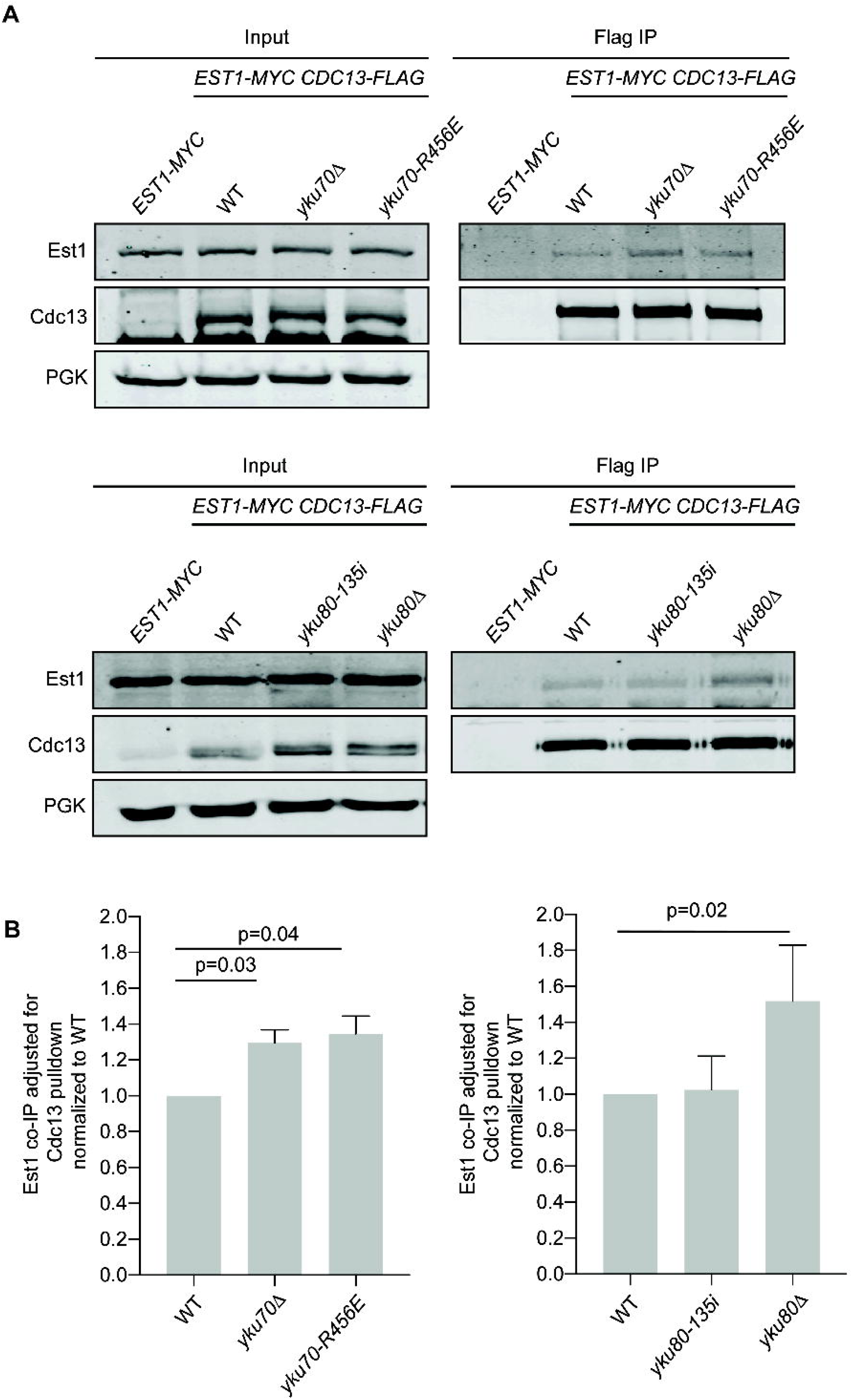
Soluble Est1:Cdc13 complexes increase in the absence of Ku. (A) Co-immunoprecipitation of Est1-myc with Cdc13-FLAG. Anti-FLAG immunoprecipitations were performed with asynchronous whole cell lysates of indicated strains. Immunoprecipitates and inputs were analyzed by western blotting with α-myc (Est1) and α-FLAG (Cdc13). Inputs were also probed with α-PGK for loading. Full-length blots are presented in Supplementary Figure S17 and S18. (B) Quantification of mean Est1-myc association relative to Cdc13-FLAG pull-down in five independent experiments for each set of genotypes. Error bars represent ± SD and p values adjusted for multiple comparisons are shown.

Levels of soluble Est1 and Cdc13 proteins were similar between strains regardless of Ku status (Supplementary Fig. S5). Thus, the increased interaction between Est1 and Cdc13 could not be attributed to potential changes in protein solubility that might have resulted from differences in the strain genotype. Notably, a similar increase in Est1:Cd13 interaction was observed in *yku70-R456E* mutant cells, indicating impaired DEB is sufficient to alter Est1:Cdc13 soluble complexes (Fig. 7A,B). Because the presence of Est1 at the telomere decreases in the absence of Ku (Fig. 6 and ref ^33^), it is likely that the increase in Est1:Cdc13 interaction observed represented complexes not associated with the telomere end. Additionally, the *yku80-135i* mutant, which maintains binding to the DNA end^31^, had levels of soluble Est:Cdc13 complexes similar to wild type (Fig. 7B,C). These findings suggest that, in the absence of Ku at the telomere, Est1 is destabilized from the end and free to bind soluble (non-telomere associated) Cdc13 resulting in an increase in Est1:Cdc13 complex formation.

## DISCUSSION

While two genome-wide screens revealed that a vast number of non-essential genes impact telomere length in *S. cerevisiae*, *YKU70* and *YKU80* fall into a rare class with respect to the markedly short telomeres observed when these genes are deleted^45, 46^. The study presented here contributes to a body of evidence that indicates that the multifunctional Ku heterodimer impacts telomere length in multiple ways, and highlights its association with the telomere end as being required and most impactful.

This work underscores the model that Ku must bind both the telomere end and TLC1 to facilitate proper telomere length maintenance^31, 44^. However, direct tethering of telomerase to the telomere by Ku is deemed unlikely based on the mutually exclusive binding of Ku to these nucleic acids^14, 43^. Whether there are two pools of Ku simultaneously functioning in telomere length maintenance, one bound to TLC1 and the other to the telomere end, or Ku shuttles between binding TLC1 and the telomere end has not been determined. Studies with recombinant Ku and Cdc13, however, indicate that Ku must load on to the telomeric end before Cdc13^47^. In addition, ChIP experiments suggest Ku is constitutively at the telomere^41^. Together, these data suggest that two pools of Ku are engaged in telomere length maintenance functions. Strikingly, we found that Ku’s interaction with Est1 was decreased when Ku’s DEB was impaired and this was not rescued by increasing TLC1 (Fig. 5A-D). This suggests that either most of Ku’s association with an Est1/TLC1 complex or the most stable form of an Est1/TLC1/Ku complex must require Ku’s prior engagement with the telomere end.

The Yku70-R456E/Yku80 DEB mutant heterodimer is proficient for Sir4 and TLC1 binding and retains TLC1 within the nucleus^33, 44^, yet telomeres were much shorter in the *yku70-R456E* mutant strain than in strains lacking Sir4, Ku:Sir4 interaction, or Ku:TLC1 interaction (Fig. 1). Additionally, TLC1 levels were decreased in the *yku70-R456E* mutant (Fig. 2), yet overexpression of TLC1 failed to rescue the short telomeres in this mutant (Fig. 3). It is important to note that we did not observe shortening of telomeres in wild type cells when TLC1 was overexpressed from the *ADH1* promoter on 2 micron plasmids (Fig. 3C) in contrast to what was observed for a modified VII-L telomere when TLC1 was overexpressed from a galactose-inducible promoter^8^ as this raises the possibility that the level of TLC1 was not sufficiently increased. However, the finding that the level of nuclear TLC1 in the *yku70-R456E* mutant was increased to a level greater than that of TLC1 in the wild type strain transformed with the empty 2 micron vector (Fig. 3D) argues that restoring nuclear TLC1 is insufficient to rescue telomere length.

Taken together, these data indicate that the short telomere phenotype of the *yku70-R456E* mutant cannot be attributed to decreased TLC1 levels. However, this is not to imply that limiting TLC1 levels are not capable of resulting in stably short telomeres in budding yeast. Mozdy et al., determined that there are ~37 molecules of TLC1 per wild type diploid^48^. This results in 0.29 molecules/chromosome end (based on 16 chromosomes × 2 in a diploid × 2 following DNA replication when telomerase acts × 2 ends/chromosome). They also determined that a *tlc1∆/TLC1* diploid has approximately half the number of TLC1 molecules/cell, at ~19, which results in 0.14 molecules per chromosome end. This amount is not sufficient to maintain normal telomere length given *tlc1∆/TLC1* heterozygous diploid cells have telomeres that are ~40 bp shorter than wild type diploids. Haploids have ~28 molecules of TLC1/cell^48^, thus 0.44 molecules/chromosome end after DNA replication. Our data indicate that there is a 50% reduction in TLC1 in a *yku70-R456E* haploid, therefore ~14 molecules/cell, resulting in ~0.22 TLC1 molecules/end after DNA replication, which is greater than that in *tlc1∆/TLC1* diploid cell (0.14), which have only 40 bp of telomere shortening. These calculations further support the conclusion that a reduction in *TLC1* is not driving the shortened telomeres in the *yku70-R456E* mutant, which are on average 161 bp shorter than wild type.

Our results suggest that the telomere length defect of the *yku70-R456E* mutant is a direct result of either impairment of Est1 recruitment or retention at the telomere. We found loss of Ku’s DEB activity results in a decrease in Est1 association with telomeric chromatin (Fig. 6). In addition, whereas tethering Est1 via Cdc13 to the telomere results in progressive and extensive elongation of telomeres in the absence of Ku’s DEB activity, tethering Est2 to the telomere does not^33^, similar to strains lacking Est1^19^. Genetic experiments have identified charge swap mutations in *EST1* and *CDC13* (*est1-60* and *cdc13-2*) that cause telomere shortening and cellular senescence when expressed individually but maintain viability and telomere length when co-expressed *in vivo*^16, 18^. A proposed model for this phenomenon is that the specific substitutions in Est1-60 and Cdc13-2 (K144E and E252K, respectively) restore a salt bridge between the proteins, facilitating telomerase recruitment to the telomere. Interestingly, the *est1-60* and *cdc13-2* mutations have little to no impact on Est1:Cdc13 interaction *in vitro*, yet the *cdc13-2* mutation impairs Est1 association with the telomere *in vivo*^14, 49^. This suggests that in the context of the cell or when Est1 is in association with telomerase, additional factors are required for stabilizing Est1’s telomere association. Our results suggest that telomere-bound Ku plays a role in this stabilization. Future studies are needed to determine how Ku influences Est1:Cdc13 interaction *in vivo*.

Like Ku and telomerase in yeast, there is evidence that Ku associates with human telomerase. Ku has been shown to independently interact with both hTR and TERT^50, 51^. Whether this binding impacts hTR levels or localization remains to be determined. Furthermore, Ku is essential in human cells because of its role in suppressing telomere deletion events^52^. In addition to directly binding telomeric DNA, human Ku also interacts with protein components of the shelterin complex - TRF1, TRF2 and Rap1^53–57^. Thus, Ku’s DEB activity may also be important for maintenance of human telomeres but further investigation is required to explore the mechanisms by which it preserves telomere length homeostasis in human cells.

## METHODS

### Yeast strains and plasmids

Strains and plasmids used in this study are described in Supplementary Table S1 and Supplementary Table S2, respectively. Gene deletions and C-terminal epitope tags were integrated using one-step allele replacement with the indicated selectable marker. Integration of mutant alleles was performed using the pop-in/pop-out technique. Strains and plasmids are available upon request.

### Protein co-immunoprecipitation assays

To analyze Est1:Cdc13 complexes, strains were grown at 28°C in 250 mL of YPD media to OD_600_=1.0. Extracts were lysed in TMG (10 mM Tris-HCl, pH 8.0, 1 mM MgCl_2_, 10% glycerol, 0.1 mM EDTA) plus 50 mM NaCl, 0.5% Tween-20, PMSF (1:10 dilution), and Set III Protease inhibitor cocktail (Calbiochem, 1:10 dilution) using acid washed glass beads. Seven milligrams of protein was incubated with 40 μL α-FLAG M2 magnetic beads (Sigma M8823) for 1 hour at 4°C. Beads were washed as follows: TMG plus 50 mM NaCl and 0.5% Tween-20 (1X); TMG plus 300 mM NaCl and 0.5% Tween-20 (3X); TMG plus 500 mM NaCl and 0.5% Tween-20 (1X); TMG plus 50 mM NaCl (1X). Immunoprecipitates and 70 μg inputs were analyzed by SDS-PAGE on 7.5% gels and transferred to an Immobilon-FL PVDF membrane (Millipore). Membranes were probed with α-FLAG (Sigma F7425, 1:1,000 dilution; Sigma F3165, 1:5,000 dilution), α-myc (Sigma C3956, 1:1,000 dilution; Sigma M4439, 1:5,000 dilution), and α-PGK (Abcam ab113687, 1:5,000 dilution) primary antibodies and quantified using ImageStudio software (LiCor). The Est1:Ku80 interaction was analyzed as above except 4 mg of protein was incubated with 30 μL α-FLAG M2 magnetic beads (Sigma M8823) and samples were analyzed by SDS-PAGE on a 10% gel.

### RNA co-immunoprecipitation assays

Experiments were performed as previously described^22^ with minor modifications. Strains were grown at 28°C in 250 mL of YPD media to OD_600_=1.0. Extracts were lysed in TMG (10 mM Tris-HCl, pH 8.0, 1 mM MgCl_2_, 10% glycerol) plus 200 mM NaCl, PMSF (1:10 dilution), Set III Protease inhibitor cocktail (Calbiochem, 1:10 dilution), and 30 μL Recombinant RNasin Ribonuclease Inhibitor (Promega). Ten milligrams of protein was incubated with 4 μL α-myc antibody (Sigma M4439) for 1 hour at 4°C, 100 μL of Pierce™ Protein G magnetic beads (Thermo Fisher Scientific 88847) was added and rotation continued for 2.5 hours. Following bead washing as previously described^22^, 10 μL of IP and 70 μg of input were reserved for western blotting. The remaining IP and input reactions were incubated at 37°C for 30 minutes in Proteinase K solution (2 mg Proteinase K, 0.01 M Tris-HCL, 0.1 M NaCl, 1% SDS, 0.01 M EDTA). RNA extraction and subsequent northern blot analysis was performed as described below. To monitor pull-down efficiency, reserve IP and input samples were analyzed by SDS-PAGE on a 10% gel, transferred to an Immobilon-FL PVDF membrane (Millipore), and probed with α-myc antibody (Sigma C3956, 1:1,000). TLC1 RNA levels and Est1 pull-down were quantified using ImageQuant software (GE Healthcare Life Sciences).

### Chromatin immunoprecipitation assays

Strains were grown in 250 mL of YPD media at 28°C to OD_600_=1.0. Cultures were crosslinked in 36.5% formaldehyde for 15 minutes at 28°C and then incubated in 2.5 M glycine for 10 minutes at room temperature with occasional shaking. Cell lysis was performed as previously described^58^. Lysates were sonicated using a Bioruptor™ UCD-200 (Diagenode) for 10 minutes on HIGH (3X). Twelve milligrams of protein was incubated with 4 μL α-myc antibody (Sigma M4439) at 4°C overnight while rotating. Simultaneously, 100 μl Pierce™ Protein G magnetic bead (Thermo Fisher Scientific 88847) slurry was pre-blocked overnight at 4°C with *E. coli* DNA and BSA (10 mg/ml). The next day pre-blocked beads were added to the IP reaction and incubated for 2 hours at 4°C. Washes were done as previously described^58^ using conventional microcentrifuge tubes. Immunoprecipitate and input samples were incubated overnight at 65°C to reverse crosslinks. One hundred micrograms Proteinase K (20 mg/ml) was added and samples were incubated overnight at 37°C. DNA was extracted using phenol:chloroform and dot blotted in duplicate onto a Zeta-Probe^®^ GT blotting membrane (Bio-Rad) according to manufacturer’s protocol. Membranes were probed with a radiolabeled telomere DNA fragment and a TyB DNA fragment. Membranes were washed, exposed to a phosphorimager screen, and quantified using ImageQuant software (GE Healthcare Life Sciences). To monitor IP efficiency, 70 μg of input and 50 μL of IP sample were analyzed by SDS-PAGE on a 10% gel, transferred to an Immobilon-FL PVDF membrane (Millipore), and probed with α-myc antibody (Sigma M4439).

### Telomere length analysis

Genomic DNA extraction and Southern blotting were performed as previously described^33^. Telomere lengths were determined using a semi-log plot generated from the distance migrated on the same agarose gel by DNA fragments in the New England Biolabs, Inc., 2-log ladder. The lengths determined for three unique biological isolates analyzed on separate Southern blots. Average lengths +/- SD are reported in Table 1.

### RNA assays

Strains were grown up in 5 mL of YPD media at 28°C overnight. RNA was harvested using the RNeasy Mini Kit (Qiagen) and concentration was measured using a NanoDrop (Thermo Fisher Scientific). For northern blotting, RNA (10-15 μg) was separated on a 1.2% agarose/1.2% formaldehyde denaturing gel. Gels were transferred to a Zeta-Probe^®^ GT blotting membrane (Bio-Rad) and probed with radiolabeled TLC1 and ACT1 fragments. Membranes were washed, exposed to a phosphorimager screen, and quantified using ImageQuant software (GE Healthcare Life Sciences). For qPCR, 1 μg of RNA was synthesized to cDNA using the Flex cDNA synthesis kit (Quanta Biosciences). Quantitative PCR was performed on a QuantFlex 6 system (Applied Biosystems) using PowerUp SYBR Green Master Mix (Applied Biosystems) and primers for actin and TLC1.

### Cytoplasmic/nuclear fractionation

Cytoplasmic and nuclear fractions of logarithmically growing yeast cells were prepared as previously described^59^ with modifications described (http://tfiib.med.harvard.edu/wiki/index.php/Yeast_Fractionation). Fifty milliliters of logarithmically growing cells (OD_600_ <1.0) were collected by centrifugation. Cell pellets were washed successively with ice cold 10 mL H2O and 10 mL SB (1 M sorbitol and 20 mM Tris-Cl pH 7.4) and stored overnight at −80°C. The pellets were thawed on ice and washed successively with 1.5 mL PSB (20 mM Tris-Cl, pH 7.4, 2 mM EDTA, 100 mM NaCl, and 10 mM β-mercaptoethanol) and 1.5 mL SB. The pellet was resuspended in 1 mL SB and spheroplasts were prepared by adding 125 μl zymolyase 20T 10 mg/ml and incubating with rotation for 60 minutes at room temperature. One milliliter SB was added and spheroplasts were collected by microcentrifugation at 2,000 RPM for 5 minutes at 4°C. The pellet was washed a second time with 1 mL SB and spheroplasts were similarly collected. The spheroplasts were resuspended in 500 μL EBX (20 mM Tris-Cl, pH 7.4, 100 mM NaCl, 0.25% Triton X-100, 15 mM β-mercaptoethanol), 0.005% phenol red, and a protease/phosphatase/RNase inhibitor cocktail of PMSF 1:100, Set III Protease Inhibitor Cocktail (Millipore) 1:100, RNasin 1:320 and ribonucleoside vanadyl complexes 1:67), and Triton X-100 added to 0.5%. After 10 minutes on ice, the resuspension was layered over 1 mL NIB (20 mM Tris-Cl, pH 7.4, 100 mM NaCl, 1.2 M sucrose, 15 mM β-mercaptoethanol, and 50 mM Na-butyrate) plus the protease/phosphatase/RNase inhibitor cocktail in a 2 mL tube and microcentrifuged at 12,000 RPM for 15 minutes at 4°C. The upper red layer was collected and analyzed as the cytoplasmic fraction. The glassy white pellet was resuspended in 500 μL EBX with Triton X-100 increased to 1% and chilled for 10 minutes on ice to yield the nuclear fraction.

For western analysis, equal volumes of each fraction were analyzed by SDS-PAGE on a 12.5% gel and transferred to Immobilon-FL PVDF membrane (Millipore). Membranes were probed with α-tubulin (Abcam ab6161; 1:1,000) primary antibody, and imaged by Odyssey Imaging System (LiCor). For northern analysis, RNA from equal volumes of each fraction was isolated using 1 mL Trizol reagent (Invitrogen), separated on a 1.2% agarose/1.2% formaldehyde denaturing gel, transferred to Zeta-Probe GT membrane (Bio-Rad), and probed with a radiolabeled TLC1 fragment.

### Statistical analyses

Statistical analyses were performed using GraphPad Prism 7.0. Data were normally distributed as determined by the Shapiro-Wilk test. Welch’s unpaired t-test was used to calculate differences when a single genotype was compared to a control. A one-way ANOVA followed by a Holm-Sidak’s multiple comparisons test was used to calculate differences when multiple genotypes were compared to a control to maintain a family error rate of <0.05. Graphs display means with error bars representing the standard deviation (SD) or standard error of the mean (SEM) as indicated. Statistical significance was assigned at p<0.05 or at adjusted p<0.05 using the Holm-Sidak’s correction for multiple comparisons if indicated in the figure legend.

### Data availability

No datasets were generated or analyzed during the current study.

## ACKNOWLEDGEMENTS

We are grateful to members of the Bertuch lab for helpful discussions. This work was supported by the National Institutes of Health, National Institute of Aging grant 5T32AG000183-23 (to L.D.L.) and National Institute of General Medical Sciences grants 5T32GM008231-22 (to L.D.L) and R01GM077509-07 (to A.A.B.).

## AUTHOR CONTRIBUTIONS

Conceived and designed the experiments: L.D.L. and A.A.B. Performed the experiments: L.D.L. and D.K.M. Analyzed data: L.D.L, D.K.M, and A.A.B. Contributed reagents/materials/analysis tools: A.A.B. Wrote the paper: L.D.L. and A.A.B.

## COMPETING INTERESTS

The authors declare no competing interests.

## Notes

#### Summary of Updates

The abstract has been revised for clarity. Figure 3D and the corresponding legend have been revised to denote the TLC1 band and a presumed nonspecific band. Supplementary Figures S7 and S10 now show RNA size markers.

## REFERENCES

1. de Lange, T. Shelterin-Mediated telomere protection. Annu Rev Genet 52, 223–247 (2018).

2. Harley, C.B., Vaziri, H., Counter, C.M. & Allsopp, R.C. The telomere hypothesis of cellular aging. Exp. Gerontol. 27, 375–382 (1992).

3. Pardue, M.L. & DeBaryshe, P.G. Drosophila telomeres: A variation on the telomerase theme. Fly (Austin) 2, 101–110 (2008).

4. Chen, J.J., Vol. 2011 (2011).

5. Greider, C.W. & Blackburn, E.H. Identification of a specific telomere terminal transferase activity in Tetrahymena extracts. Cell 43, 405–413 (1985).

6. Wellinger, R.J. & Zakian, V.A. Everything you ever wanted to know about Saccharomyces cerevisiae telomeres: beginning to end. Genetics 191, 1073–1105 (2012).

7. Kupiec, M. Biology of telomeres: lessons from budding yeast. FEMS Microbiol. Rev. 38, 144–171 (2014).

8. Singer, M.S. & Gottschling, D.E. TLC1: template RNA component of Saccharomyces cerevisiae telomerase. Science 266, 404–409 (1994).

9. Cohn, M. & Blackburn, E.H. Telomerase in yeast. Science 269, 396–400 (1995).

10. Lingner, J. et al. Reverse transcriptase motifs in the catalytic subunit of telomerase. Science 276, 561–567 (1997).

11. Seto, A.G., Livengood, A.J., Tzfati, Y., Blackburn, E.H. & Cech, T.R. A bulged stem tethers Est1p to telomerase RNA in budding yeast. Genes Dev. 16, 2800–2812 (2002).

12. Tucey, T.M. & Lundblad, V. Regulated assembly and disassembly of the yeast telomerase quaternary complex. Genes Dev. 28, 2077–2089 (2014).

13. Hughes, T.R., Evans, S.K., Weilbaecher, R.G. & Lundblad, V. The Est3 protein is a subunit of yeast telomerase. Curr. Biol. 10, 809–812 (2000b).

14. Chen, H. et al. Structural Insights into Yeast Telomerase Recruitment to Telomeres. Cell 172, 331–343 e313 (2018).

15. Bianchi, A., Negrini, S. & Shore, D. Delivery of yeast telomerase to a DNA break depends on the recruitment functions of Cdc13 and Est1. Mol. Cell 16, 139–146 (2004).

16. Pennock, E., Buckley, K. & Lundblad, V. Cdc13 delivers separate complexes to the telomere for end protection and replication. Cell 104, 387–396 (2001).

17. Qi, H. & Zakian, V.A. The Saccharomyces telomere-binding protein Cdc13p interacts with both the catalytic subunit of DNA polymerase alpha and the telomerase-associated Est1 protein. Genes Dev. 14, 1777–1788 (2000).

18. Nugent, C.I., Hughes, T.R., Lue, N.F. & Lundblad, V. Cdc13p: A single-strand telomeric DNA-binding protein with a dual role in yeast telomere maintenance. Science 274, 249–252 (1996).

19. Evans, S.K. & Lundblad, V. Est1 and Cdc13 as comediators of telomerase access. Science 286, 117–120 (1999).

20. Evans, S.K. & Lundblad, V. The Est1 subunit of Saccharomyces cerevisiae telomerase makes multiple contributions to telomere length maintenance. Genetics 162, 1101–1115 (2002).

21. Tuzon, C.T., Wu, Y., Chan, A. & Zakian, V.A. The Saccharomyces cerevisiae Telomerase Subunit Est3 Binds Telomeres in a Cell Cycle- and Est1-Dependent Manner and Interacts Directly with Est1 In Vitro. PLoS Genet. 7, e1002060 (2011).

22. Lingner, J., Cech, T.R., Hughes, T.R. & Lundblad, V. Three Ever Shorter Telomere (*EST*) genes are dispensable for in vitro yeast telomerase activity. Proc. Natl. Acad. Sci. USA 94, 11190–11195 (1997).

23. Lendvay, T.S., Morris, D.K., Sah, J., Balasubramanian, B. & Lundblad, V. Senescence mutants of *Saccharomyces cerevisiae* with a defect in telomere replication identify three additional *EST* genes. Genetics 144, 1399–1412 (1996).

24. Lundblad, V. & Szostak, J.W. A mutant with a defect in telomere elongation leads to senescence in yeast. Cell 57, 633–643 (1989).

25. Walker, J.R., Corpina, R.A. & Goldberg, J. Structure of the Ku heterodimer bound to DNA and its implications for double-strand break repair. Nature 412, 607–614 (2001).

26. Emerson, C.H. & Bertuch, A.A. Consider the workhorse: Nonhomologous end-joining in budding yeast. Biochem. Cell Biol. 94, 396–406 (2016).

27. Gravel, S., Larrivee, M., Labrecque, P. & Wellinger, R.J. Yeast Ku as a regulator of chromosomal DNA end structure. Science 280, 741–745 (1998).

28. Porter, S.E., Greenwell, P.W., Ritchie, K.B. & Petes, T.D. The DNA-binding protein Hdf1p (a putative Ku homologue) is required for maintaining normal telomere length in *Saccharomyces cerevisiae*. Nucleic Acids Res. 24, 582–585 (1996).

29. Boulton, S.J. & Jackson, S.P. Identification of a *Saccharomyces cerevisiae* Ku80 homologue: roles in DNA double strand break rejoining and in telomeric maintenance. Nucleic Acids Res. 24, 4639–4648 (1996).

30. Peterson, S.E. et al. The function of a stem-loop in telomerase RNA is linked to the DNA repair protein Ku. Nat. Genet. 27, 64–67 (2001).

31. Stellwagen, A.E., Haimberger, Z.W., Veatch, J.R. & Gottschling, D.E. Ku interacts with telomerase RNA to promote telomere addition at native and broken chromosome ends. Genes Dev. 17, 2384–2395 (2003).

32. Gallardo, F., Olivier, C., Dandjinou, A.T., Wellinger, R.J. & Chartrand, P. TLC1 RNA nucleo-cytoplasmic trafficking links telomerase biogenesis to its recruitment to telomeres. EMBO J. 27, 748–757 (2008).

33. Williams, J.M., Ouenzar, F., Lemon, L.D., Chartrand, P. & Bertuch, A.A. The principal role of ku in telomere length maintenance is promotion of est1 association with telomeres. Genetics 197, 1123–1136 (2014).

34. Mozdy, A.D., Podell, E.R. & Cech, T.R. Multiple yeast genes, including Paf1 complex genes, affect telomere length via telomerase RNA abundance. Mol. Cell. Biol. 28, 4152–4161 (2008).

35. Zappulla, D.C. et al. Ku can contribute to telomere lengthening in yeast at multiple positions in the telomerase RNP. RNA 17, 298–311 (2011).

36. Moretti, P., Freeman, K., Coodly, L. & Shore, D. Evidence that a complex of SIR proteins interacts with the silencer and telomere-binding protein RAP1. Genes Dev. 8, 2257–2269 (1994).

37. Gottschling, D.E., Aparicio, O.M., Billington, B.L. & Zakian, V.A. Position effect at S. cerevisiae telomeres: reversible repression of Pol II transcription. Cell 63, 751–762 (1990).

38. Hass, E.P. & Zappulla, D.C. The Ku subunit of telomerase binds Sir4 to recruit telomerase to lengthen telomeres in S. cerevisiae. Elife 4 (2015).

39. Bertuch, A.A. & Lundblad, V. The Ku heterodimer performs separable activities at double strand breaks and chromosome termini. Mol. Cell. Biol. 23, 8202–8215 (2003).

40. Ribes-Zamora, A., Mihalek, I., Lichtarge, O. & Bertuch, A.A. Distinct faces of the Ku heterodimer mediate DNA repair and telomeric functions. Nat. Struct. Mol. Biol. 14, 301–307 (2007).

41. Fisher, T.S., Taggart, A.K. & Zakian, V.A. Cell cycle-dependent regulation of yeast telomerase by Ku. Nat. Struct. Mol. Biol. 11, 1198–1205 (2004).

42. Chan, A., Boule, J.B. & Zakian, V.A. Two pathways recruit telomerase to Saccharomyces cerevisiae telomeres. PLoS Genet. 4, e1000236 (2008).

43. Pfingsten, J.S. et al. Mutually exclusive binding of telomerase RNA and DNA by Ku alters telomerase recruitment model. Cell 148, 922–932 (2012).

44. Lopez, C.R. et al. Ku must load directly onto the chromosome end in order to mediate its telomeric functions. PLoS Genet. 7, e1002233 (2011).

45. Askree, S.H. et al. A genome-wide screen for Saccharomyces cerevisiae deletion mutants that affect telomere length. Proc. Natl. Acad. Sci. U S A 101, 8658–8663 (2004).

46. Gatbonton, T. et al. Telomere length as a quantitative trait: genome-wide survey and genetic mapping of telomere length-control genes in yeast. PLoS Genet. 2, e35 (2006).

47. Wu, T.J. et al. Sequential loading of Saccharomyces cerevisiae Ku and Cdc13p to telomeres. J. Biol. Chem. 284, 12801–12808 (2009).

48. Mozdy, A.D. & Cech, T.R. Low abundance of telomerase in yeast: implications for telomerase haploinsufficiency. RNA 12, 1721–1737 (2006).

49. Wu, Y. & Zakian, V.A. The telomeric Cdc13 protein interacts directly with the telomerase subunit Est1 to bring it to telomeric DNA ends in vitro. Proc. Natl. Acad. Sci. U S A 108, 20362–20369 (2011).

50. Ting, N.S., Yu, Y., Pohorelic, B., Lees-Miller, S.P. & Beattie, T.L. Human Ku70/80 interacts directly with hTR, the RNA component of human telomerase. Nucleic Acids Res. 33, 2090–2098 (2005).

51. Chai, W., Ford, L.P., Lenertz, L., Wright, W.E. & Shay, J.W. Human Ku70/80 associates physically with telomerase through interaction with hTERT. J. Biol. Chem. 277, 47242–47247 (2002).

52. Wang, Y., Ghosh, G. & Hendrickson, E.A. Ku86 represses lethal telomere deletion events in human somatic cells. Proc. Natl. Acad. Sci. U S A 106, 12430–12435 (2009).

53. Hsu, H.L. et al. Ku acts in a unique way at the mammalian telomere to prevent end joining. Genes Dev. 14, 2807–2812 (2000).

54. Hsu, H.L., Gilley, D., Blackburn, E.H. & Chen, D.J. Ku is associated with the telomere in mammals. Proc. Natl. Acad. Sci. U S A 96, 12454–12458 (1999).

55. O’Connor, M.S., Safari, A., Liu, D., Qin, J. & Songyang, Z. The human Rap1 protein complex and modulation of telomere length. J. Biol. Chem. 279, 28585–28591. Epub 22004 Apr 28520. (2004).

56. Song, K., Jung, D., Jung, Y., Lee, S.G. & Lee, I. Interaction of human Ku70 with TRF2. FEBS Lett. 481, 81–85 (2000).

57. Ribes-Zamora, A., Indiviglio, S.M., Mihalek, I., Williams, C.L. & Bertuch, A.A. TRF2 interaction with Ku heterotetramerization interface gives insight into c-NHEJ prevention at human telomeres. Cell Rep. 5, 194–206 (2013).

58. Aparicio, O.M. et al. Chromatin immunoprecipitation for determining the association of proteins with specific genomic regions in vivo, in Current Protocols in Molecular Biology. (eds. F.M. Ausubel et al.) 21.23.21–21.23.17 (John Wiley & Sons, Inc., 2005).

59. Keogh, M.C. et al. The Saccharomyces cerevisiae histone H2A variant Htz1 is acetylated by NuA4. Genes Dev. 20, 660–665 (2006).

