## Supplementary material for "Loss of Ku’s DNA end binding activity affects telomere length via destabilizing telomere-bound Est1 rather than altering TLC1 homeostasis": Supplentary Information and Figures S1-S18

Alison A. Bertuch, MD, PhD

1102 Bates, FC 1200

Houston, TX, 77030

**Supplementary Table S1.** Strains used in the study

| STRAIN | GENOTYPE | SOURCE |
|---|---|---|
| YAB289 | *MAT**a** ura3-52 lys2-801 ade2-101 trp-Δ1 his3-Δ200 leu2-Δ1* | [1] |
| YAB200 | YAB289 *yku70Δ::KAN$^R$* | This study |
| YAB470 | YAB289 *cdc13Δ::NAT$^R$ yku70Δ::HPH$^R$* pVL438 | [1] |
| YAB471 | YAB289 *cdc13Δ::NAT$^R$* pVL438 | [1] |
| YAB540 | YAB289 *yku70-R456E* | This study |
| YAB620 | YAB289 *cdc13Δ::NAT$^R$ yku70-R456E* pVL438 | [1] |
| YAB621 | YAB289 *yku80-135i* | [1] |
| YAB766 | YAB289 *yku80Δ::NAT$^R$* | [1] |
| YAB852 | YAB289 *tlc1Δ::KAN$^R$* pSD120 | This study |
| YAB928 | YAB289 *CDC13-G6-(FLAG)$_3$::KAN$^R$ EST1-(MYC)$_{13}$::HIS3 yku80Δ::HPH$^R$* | This study |
| YAB930 | YAB289 *CDC13-G6-(FLAG)$_3$::KAN$^R$ EST1-(MYC)$_{13}$::HIS3* | This study |
| YAB936 | YAB289 *EST1-(MYC)$_{13}$::HIS3* | This study |
| YAB937 | YAB289 *CDC13-G6-(FLAG)$_3$::KAN$^R$ EST1-(MYC)$_{13}$::HIS3 yku80-135i* | This study |
| YAB958 | YAB289 *CDC13-G6-(FLAG)$_3$::KAN$^R$ EST1-(MYC)$_{13}$::HIS3 yku70Δ::KAN$^R$* | This study |
| YAB959 | YAB289 *EST1-(MYC)$_{13}$::HIS3 yku70-R456E* | This study |
| YAB961 | YAB289 *CDC13-G6-(FLAG)$_3$::KAN$^R$ EST1-(MYC)$_{13}$::HIS3 yku70-R456E* | This study |
| YAB1021 | YAB289 *sir4Δ::KAN$^R$* | This study |
| YAB1023 | YAB289 *tlc1Δ48* | This study |
| YAB1024 | YAB289 *yku80-L111R* | This study |
| YAB1025 | YAB289 *yku80-L115A* | This study |
| YAB1027 | YAB289 *EST1-(MYC)$_{13}$::HIS3 YKU80-(FLAG)$_3$::KAN$^R$* | This study |
| YAB1028 | YAB289 *EST1-(MYC)$_{13}$::HIS3 YKU80-(FLAG)$_3$::KAN$^R$ yku70-R456E* | This study |

**Supplementary Table S2.** Plasmids used in this study

| **Plasmid** | **Genotype** | **Source** |
|---|---|---|
| pAB198 | *CEN TRP1 yku70-R456E* | [2] |
| pAB830 | *2μ URA3 ADH1-TLC1* | [1] |
| pAB889 | *CEN URA3 yku80-135i* | This study |
| pRS414 | *CEN TRP1* | [3] |
| pRS416 | *CEN URA3* | [3] |
| pRS426 | *2μ URA3* | [3] |
| pSD120 | *CEN URA3 TLC1* | [4] |
| pVL438 | *CEN URA3 CDC13* | [5] |
| pVL648 | *CEN LEU2 CDC13* | [5] |
| pVL1057 | *CEN TRP1 YKU70* | [6] |
| pVL1069 | *CEN URA3 YKU80* | [7] |
| pVL1091 | *CEN LEU2 CDC13-EST1* | [5] |

### References


1. Williams, J.M., et al., *The principal role of Ku in telomere length maintenance is promotion of Est1 association with telomeres.* Genetics, 2014. **197**(4):1123-36.
2. Lopez, C.R., et al., *Ku must load directly onto the chromosome end in order to mediate its telomeric functions.* PLoS Genet, 2011. **7**(8):e1002233.
3. Christianson, T.W., et al., *Multifunctional yeast high-copy-number shuttle vectors.* Gene 1992. **110**(1):119-22.
4. Diede, S. J., and D.E. Gottschling, *Telomerase-mediated telomere addition in vivo requires DNA primase and DNA polymerases alpha and delta.* Cell, 1999. **99**(7):723-33.
5. Evans, S.K. and V. Lundblad, Est1 and Cdc13 as comediators of telomerase access. Science,1999. **286**(5437):117-20.

6. Ribes-Zamora, A., et al., *Distinct faces of the Ku heterodimer mediate DNA repair and telomeric functions.* Nat Struct Mol Biol, 2007. **14**(4):301-7.
7. Bertuch, A.A. and V. Lundblad, *The Ku heterodimer performs separable activities at double-strand breaks and chromosome termini.* Mol Cell Biol, 2003. **23**(22):8202-15.

**Supplementary Methods**

**Protein solubility experiments**

Strains were grown at 28°C in 50 mL of YPD media to $OD_{600}$=0.8. Extracts were lysed in TMG (10 mM Tris-HCl, pH 8.0, 1 mM $MgCl_2$, 10% glycerol, 0.1 mM EDTA) plus 50 mM NaCl, PMSF (1:10 dilution), and Set III Protease inhibitor cocktail (Calbiochem, Millipore, 1:10 dilution) using acid washed glass beads. The supernatant (soluble fraction) was collected by centrifuging at 14000 RPM for 15 minutes at 4°C. The remaining cell debris pellet was resuspended in a solubilization buffer (20mM $NaPO_4$ buffer pH 8.0, 300 mM NaCl, 2% SDS, 2 mM DTT, 1% Triton X-100), PMSF (1:10 dilution), and Set III Protease inhibitor cocktail (Calbiochem, Millipore, 1:10 dilution) and the supernatant (insoluble fraction) was collected by centrifuging at 14000 RPM for 15 minutes at 4°C. Equal volume of soluble and insoluble fraction was run on a 7.5% polyacrylamide gel, transferred to an Immobilon-FL PVDF membrane (Millipore), and probed with $\alpha$-FLAG (Sigma F7425, 1:1000 dilution), $\alpha$-myc (Sigma M4439, 1:5000 dilution), and $\alpha$-PGK (Abcam ab113687, 1:5000 dilution) primary antibodies. Percent Cdc13 and Est1 was calculated per fraction.

Figure-S1 (Bertuch)

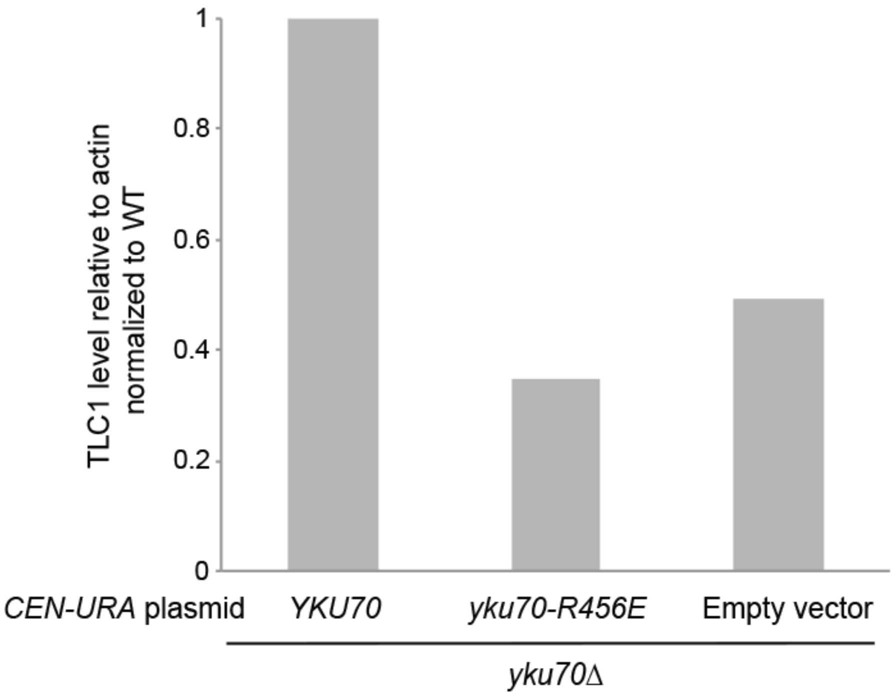


**Supplementary Figure S1. *YKU70* restores TLC1 levels in a *yku70*Δ strain while the *yku70-R456E* mutant fails to rescue TLC1 levels.** Quantification of TLC1 by RT-qPCR in asynchronous *yku70*Δ cells tranformed with indicated plasmids. TLC1 levels were quantified relative to actin RNA and normalized to levels obtained for the *YKU70 CEN-URA* transformed (WT) cells.

Figure-S2 (Bertuch)

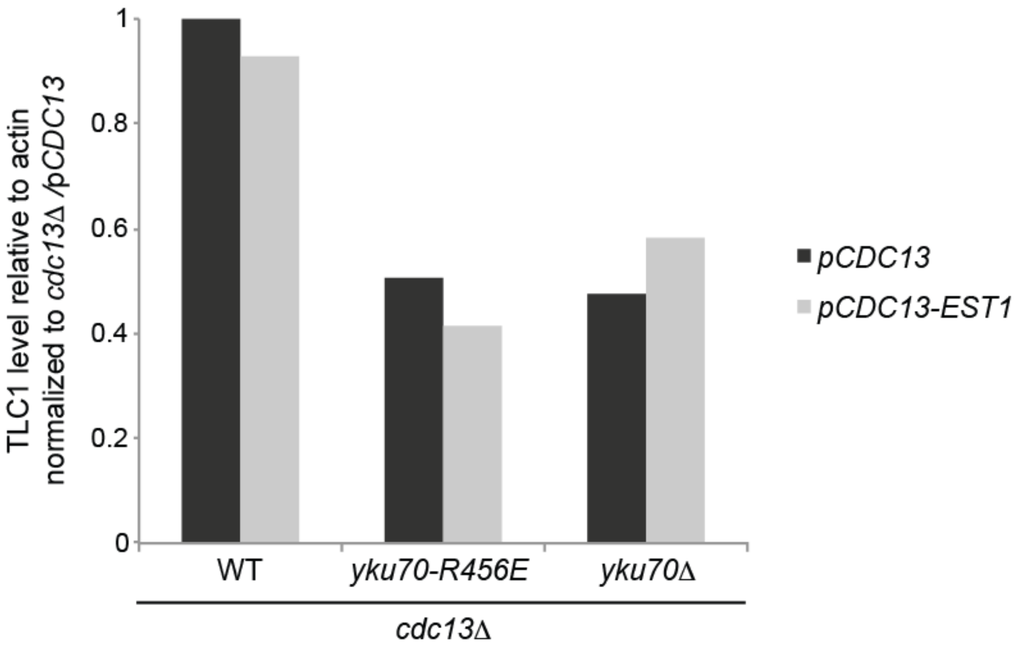


**Supplementary Figure S2. TLC1 levels are not stabilized in Ku mutant strains expressing a *CDC13-EST1* fusion.** RT-qPCR of TLC1 in asynchronous *cdc13Δ YKU70* (WT), *cdc13Δ yku70-R456E* and *cdc13Δ yku70Δ* strains expressing plasmid-borne *CDC13* or *CDC13-EST1* fusion.

Figure-S3 (Bertuch)

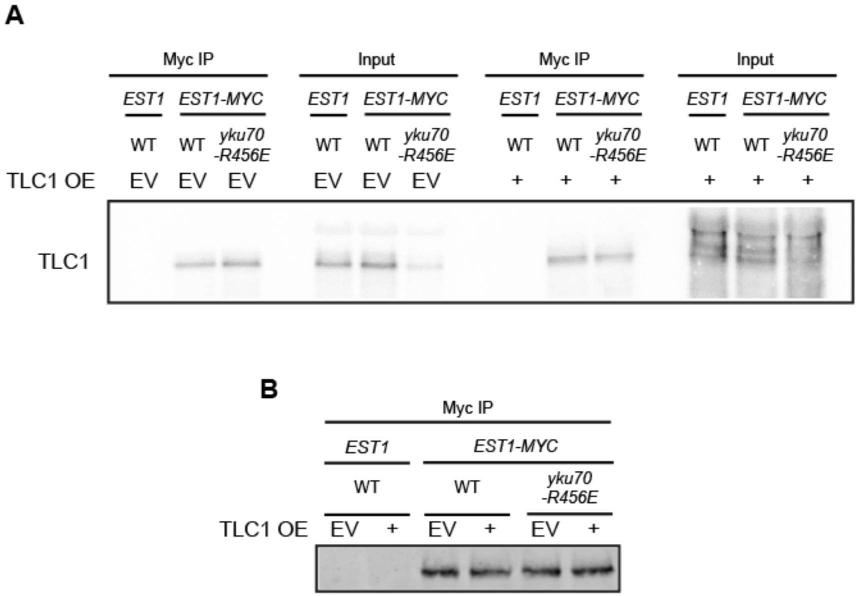


**Supplementary Figure S3. Est1-TLC1 interaction is not reduced in a *yku70-R456E* strain, with or without TLC1 overexpression.** Whole cell lystates from asychronous cultures of strains with EST1 or EST1-MYC, YKU70 (WT) or yku70-R456E, and carrying either EV or TLC1 OE 2 micron plasmids, were immunoprecipitated with anti-myc and examined by northern blot for co-IP of TLC1 (A) or western blot to detect Est1-myc (B). Full-length blots are presented in Supplementary Figure S15.

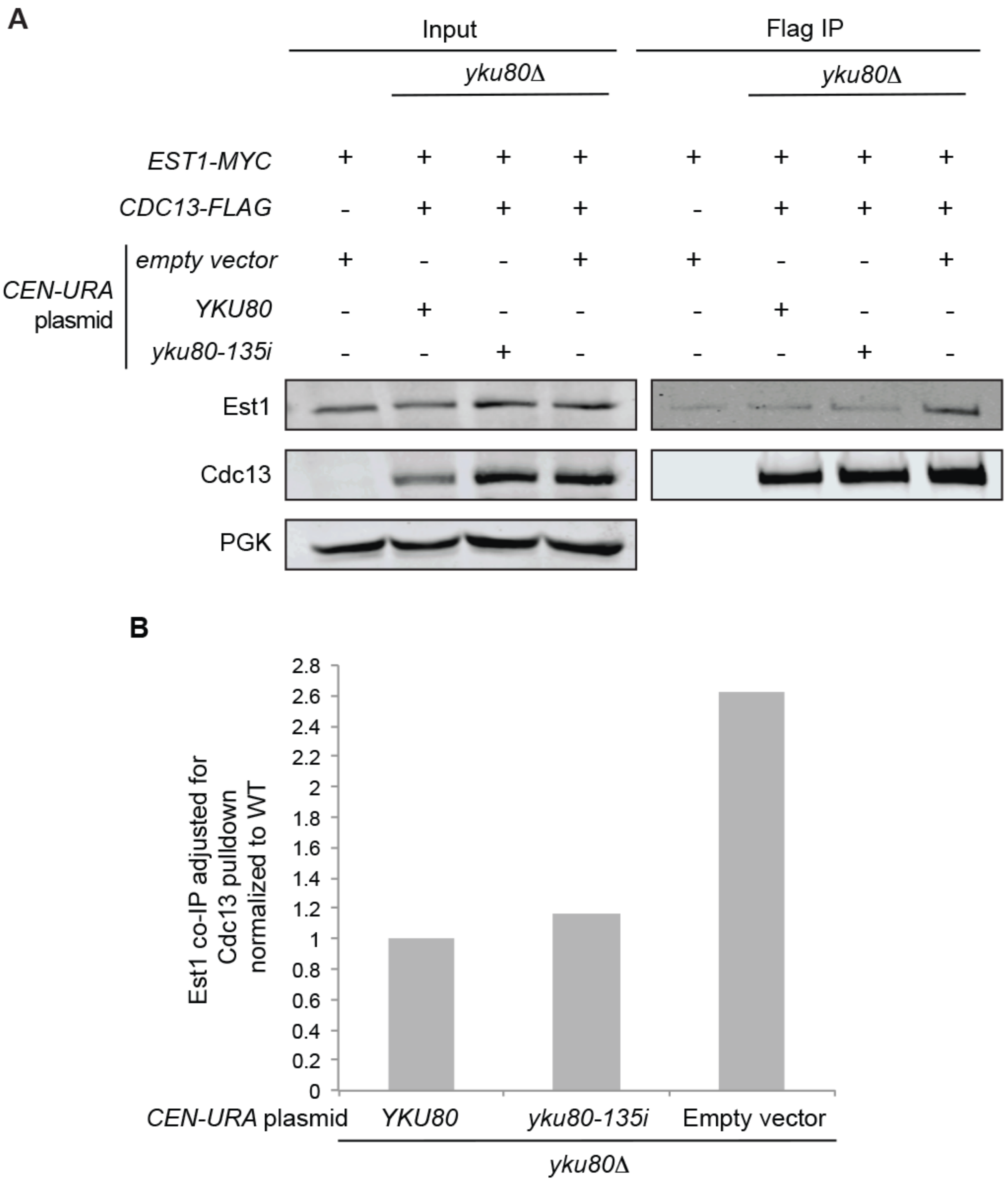


**Supplementary Figure S4. The Est1:Cdc13 interaction is rescued with *YKU80* plasmids.** (A) Co-immunoprecipitation of Est1-myc with Cdc13-FLAG in *yku80Δ* strains supplemented with indicated plasmids. Anti-FLAG immunoprecipitations were performed with whole cell lysate of asynchronous strains. Immunoprecipites and inputs were analyzed by western blotting with α-myc (Est1) and α-FLAG (Cdc13). Inputs were also probed with α-PGK for loading. Full length blots are presented in Supplementary Figure S14. (B) Quantification of Est1 associations relative to Cdc13 pull-down.

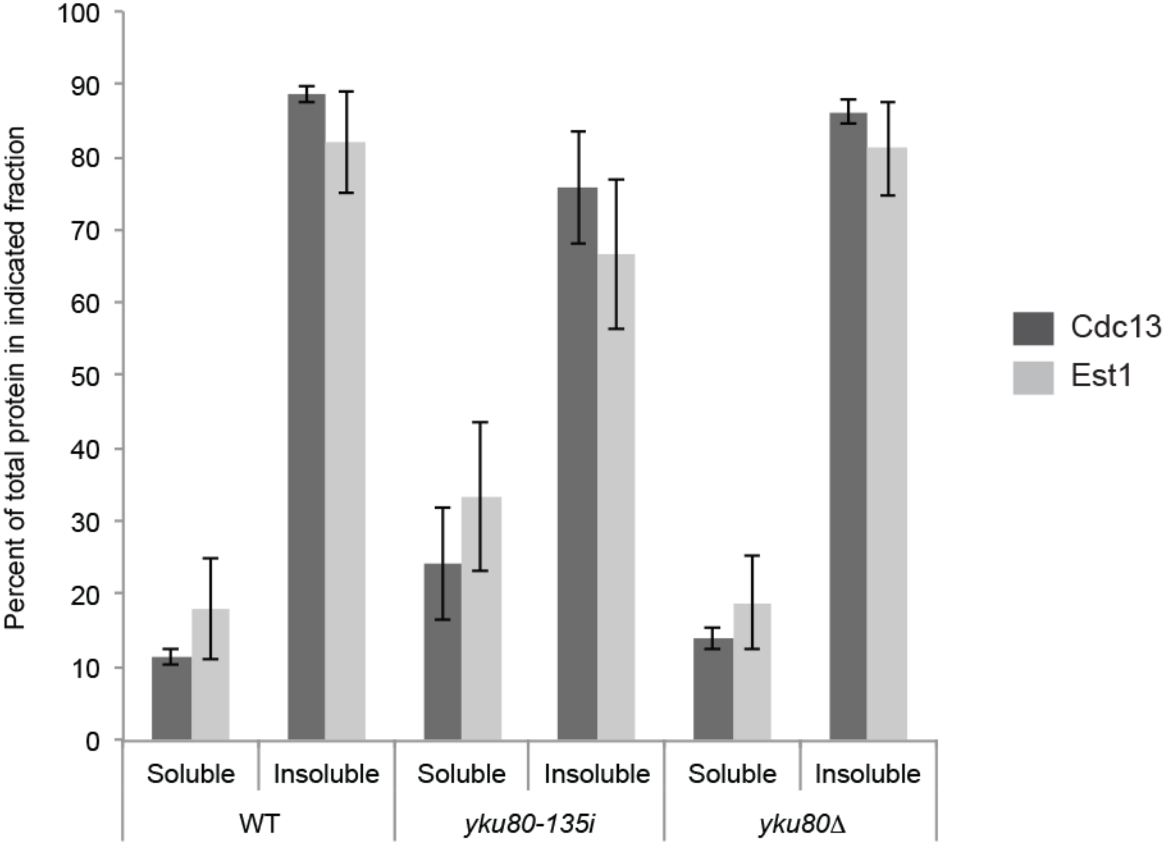


**Supplementary Figure S5. The increased Est1:Cdc13 interaction is not caused by changes in protein solubility.** Quantification of western blots for total Est1 and Cdc13 protein in soluble and insoluble fractions in three independent experiments. Indicated fractions were analyzed by western blotting with α-myc (Est1) and α-FLAG (Cdc13). Error bars represent ± 1 SEM.

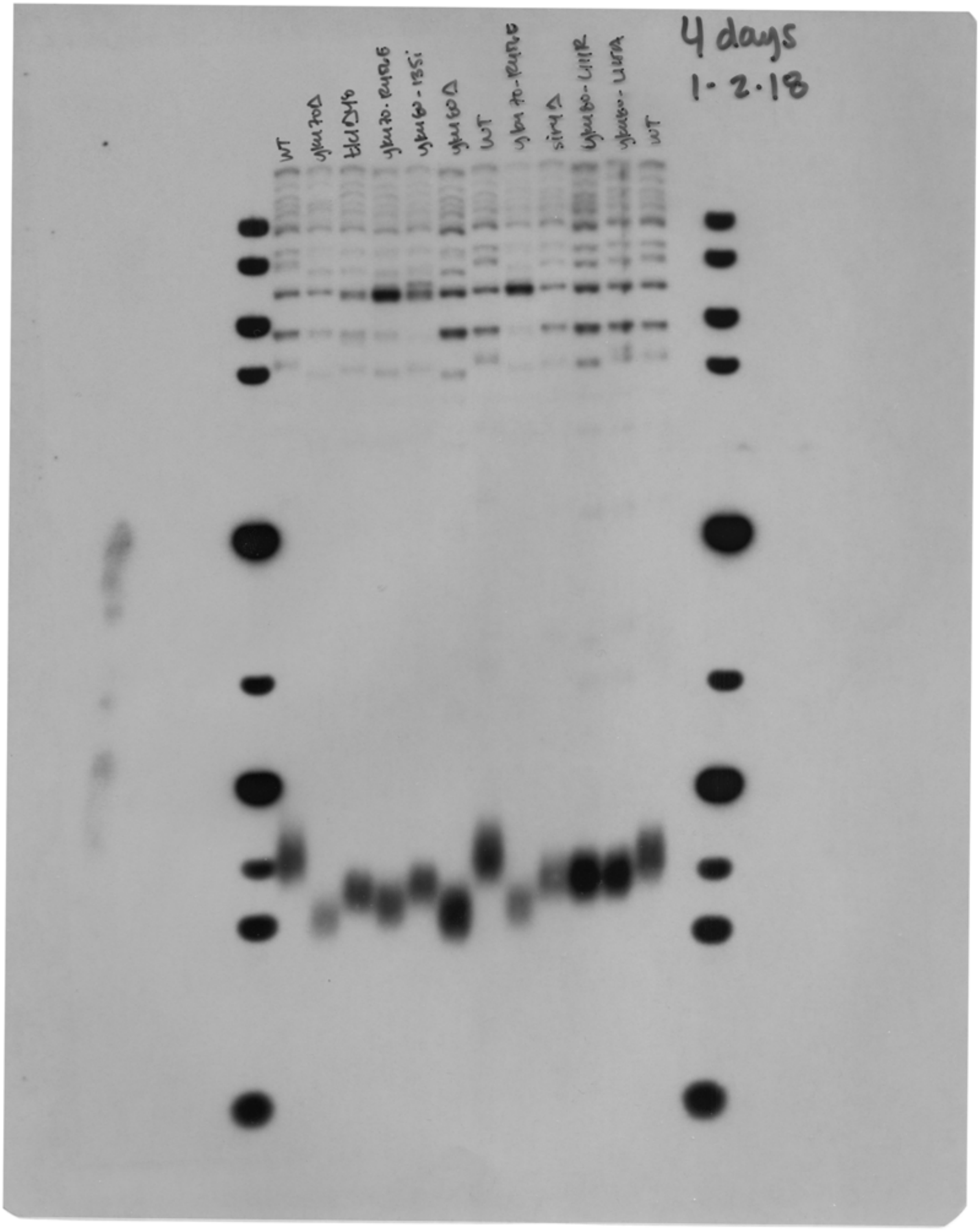


**Supplementary Figure S6.** Uncropped Southern blot shown in **Figure 1.**

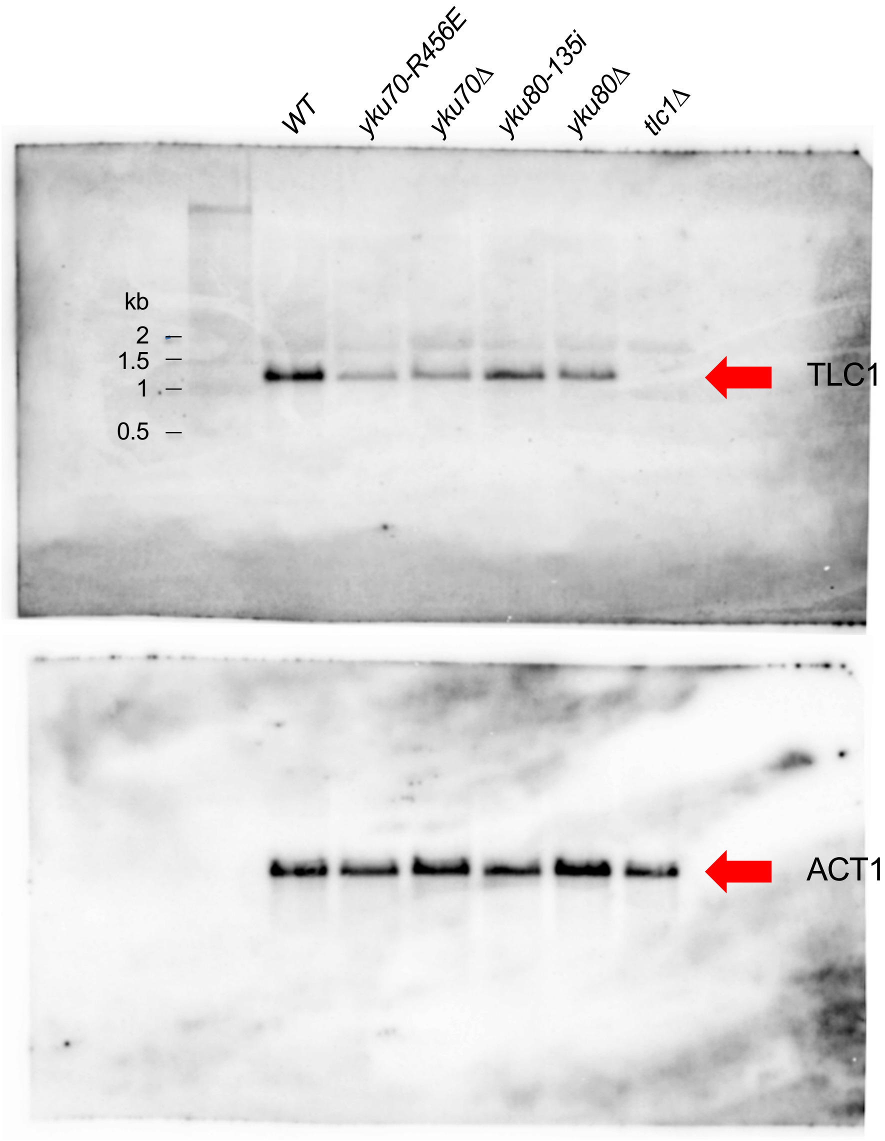


**Supplementary Figure S7.** Uncropped northern blots shown in **Figure 2B.**

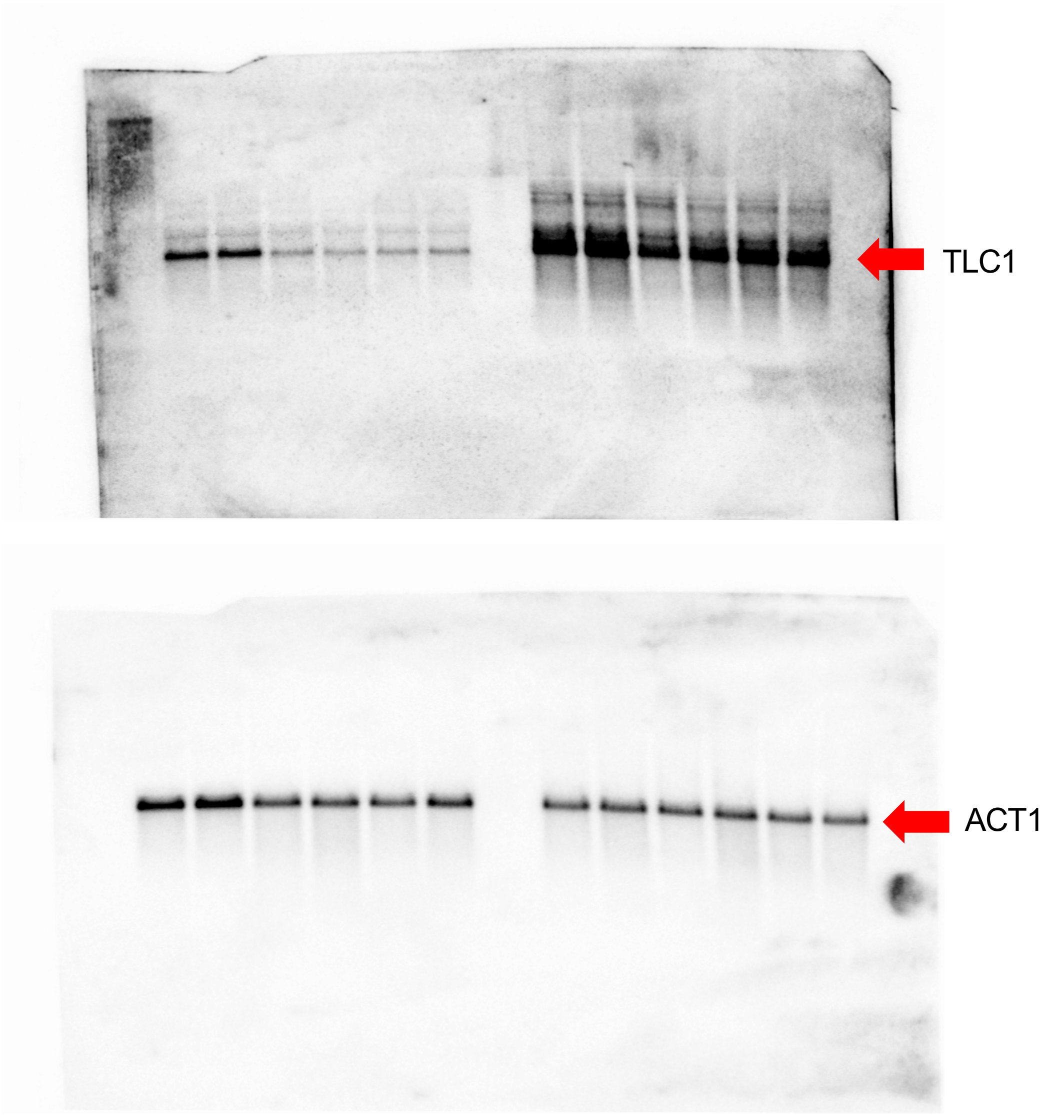


**Supplementary Figure S8.** Uncropped northern blots shown in **Figure 3A.**

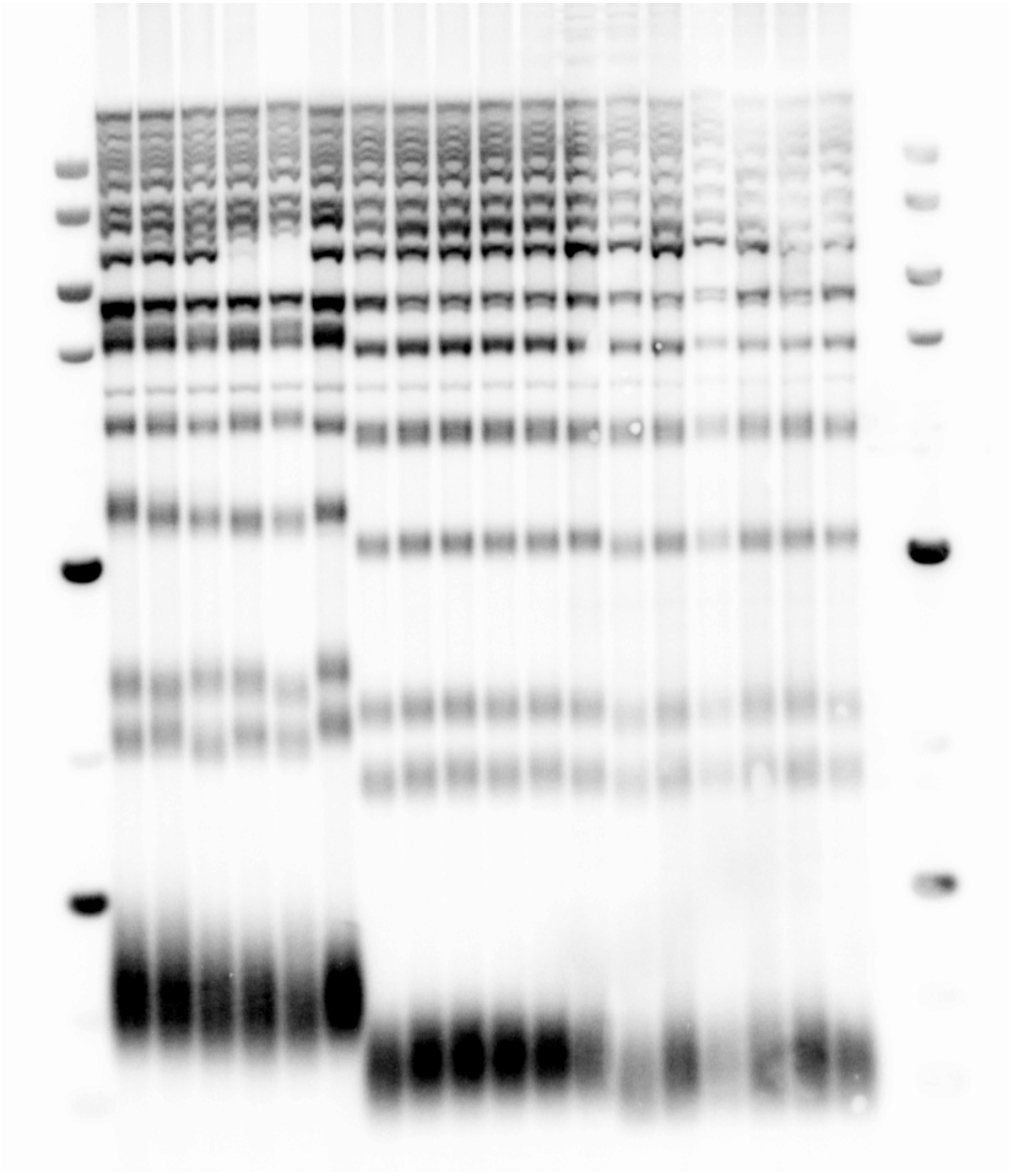

**Supplementary Figure S9.** Uncropped Southern blot shown in **Figure 3C.**

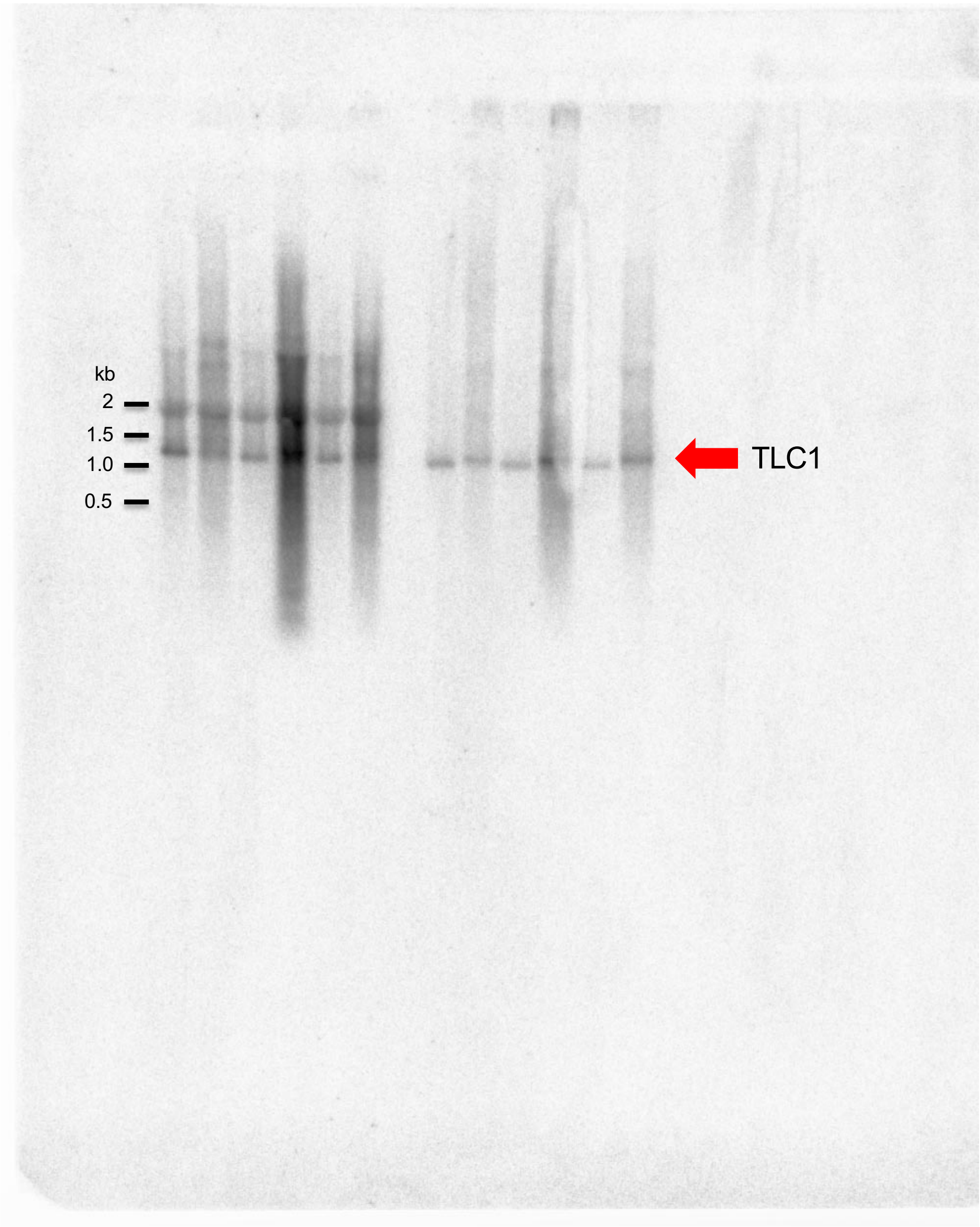


**Supplementary Figure S10A.** Uncropped TLC1 northern blot shown in **Figure 3D.**

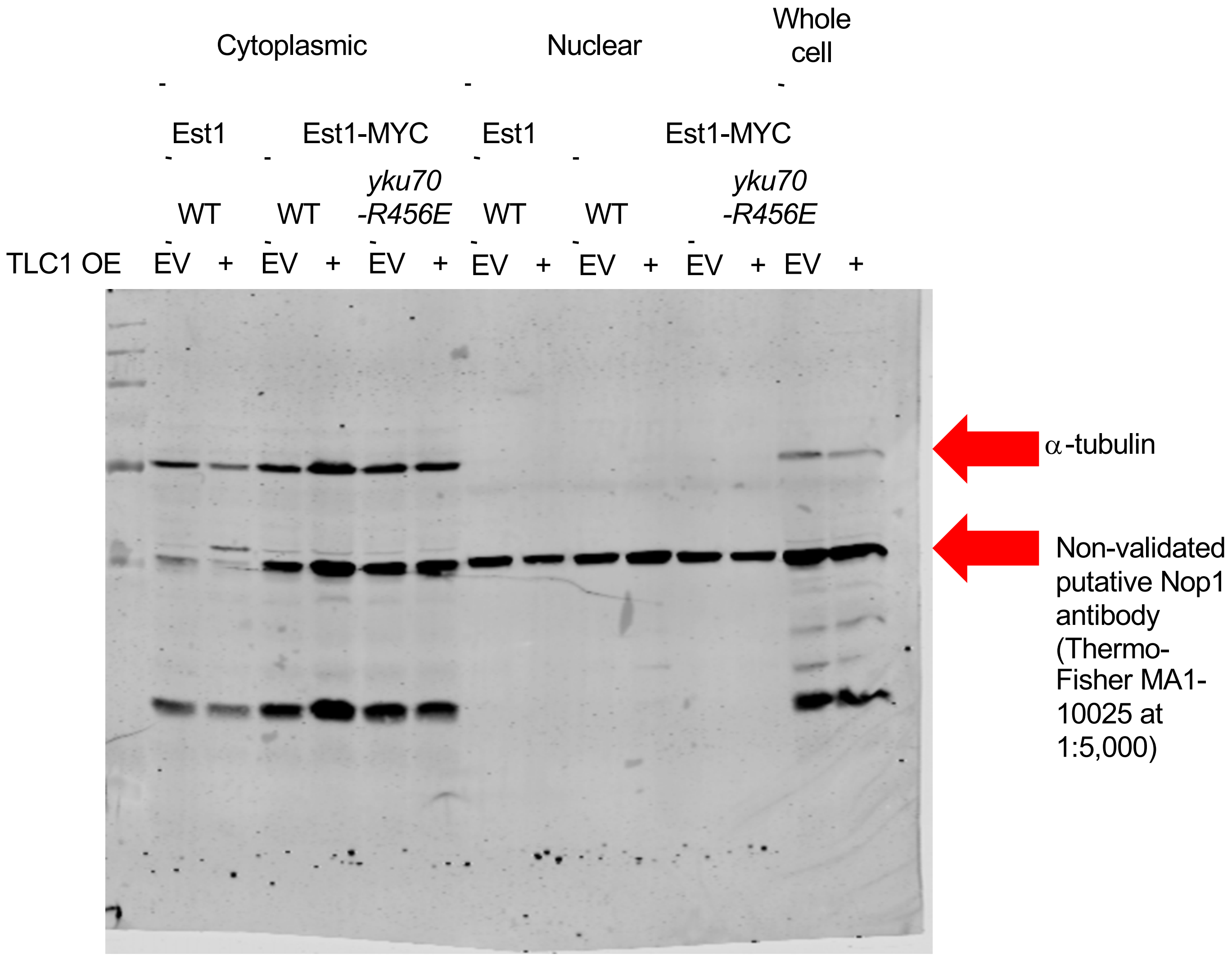


**Supplementary Figure S10B.** Uncropped western blot shown in **Figure 3D.**

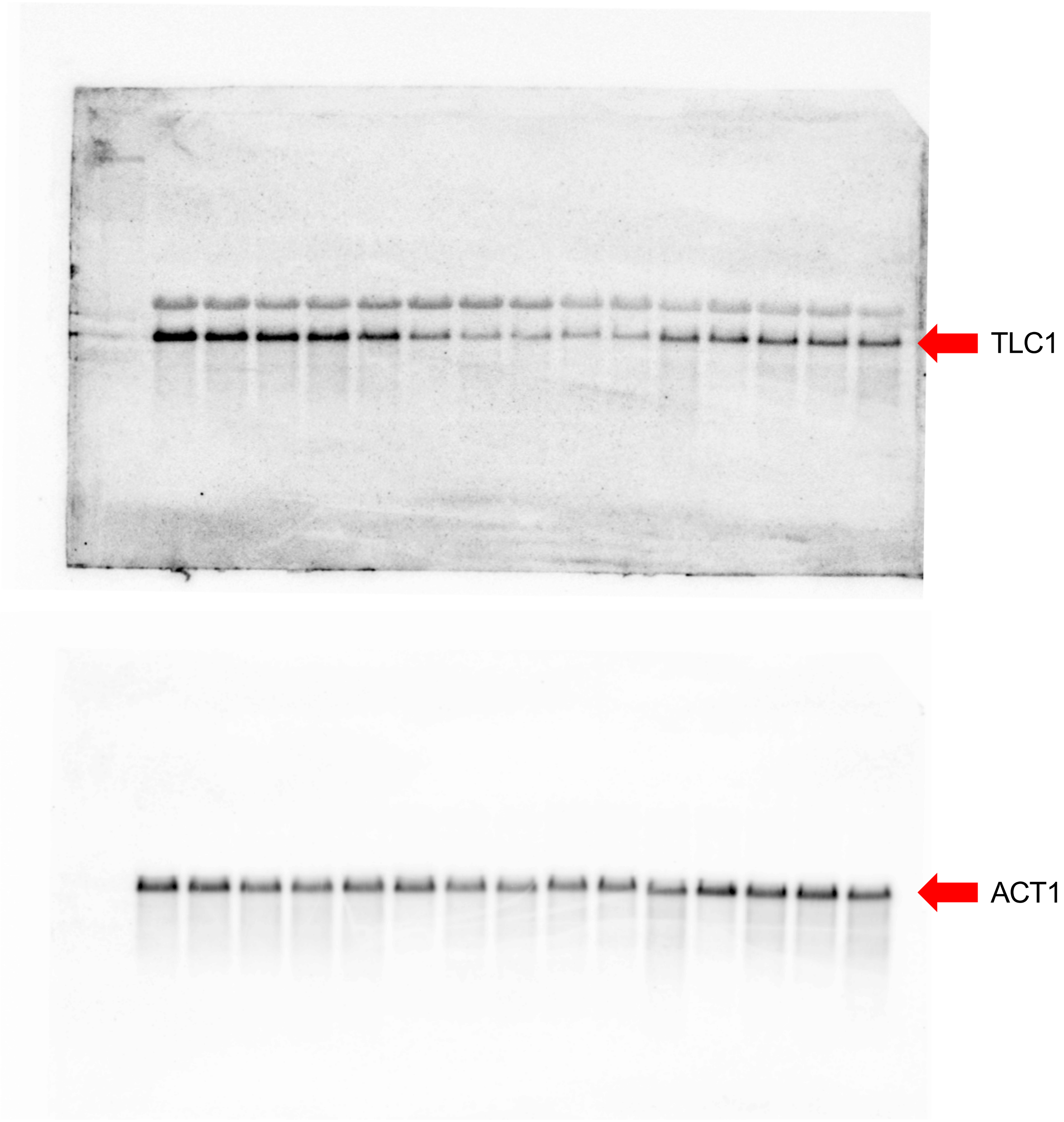


**Supplementary Figure S11.** Uncropped northern blots shown in **Figure 4A.**

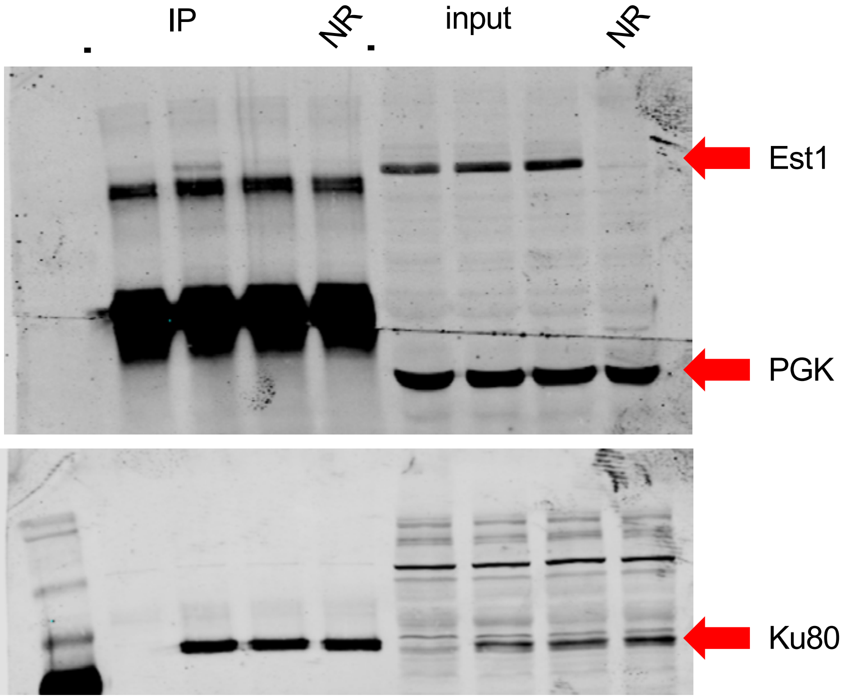


**Supplementary Figure S12.** Uncropped western blots shown in **Figure 5A**; NR indicates not relevant

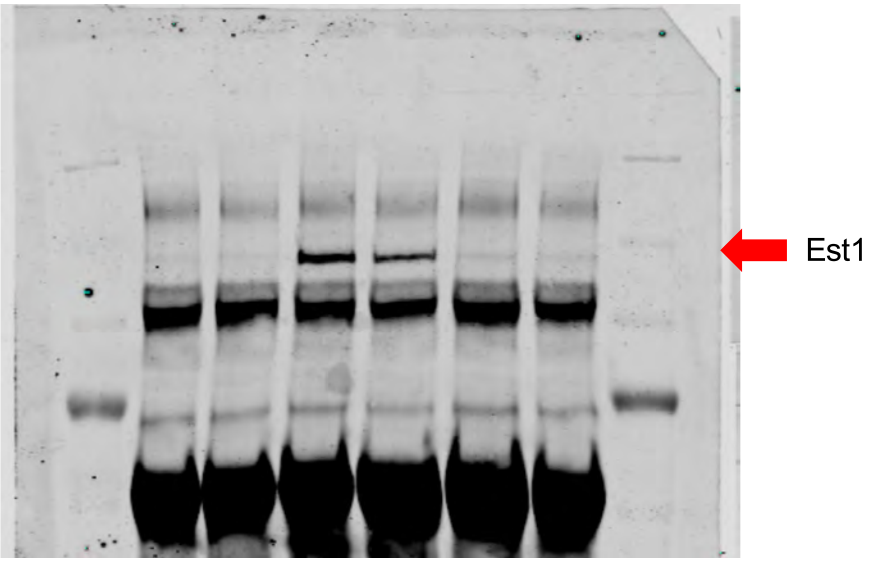


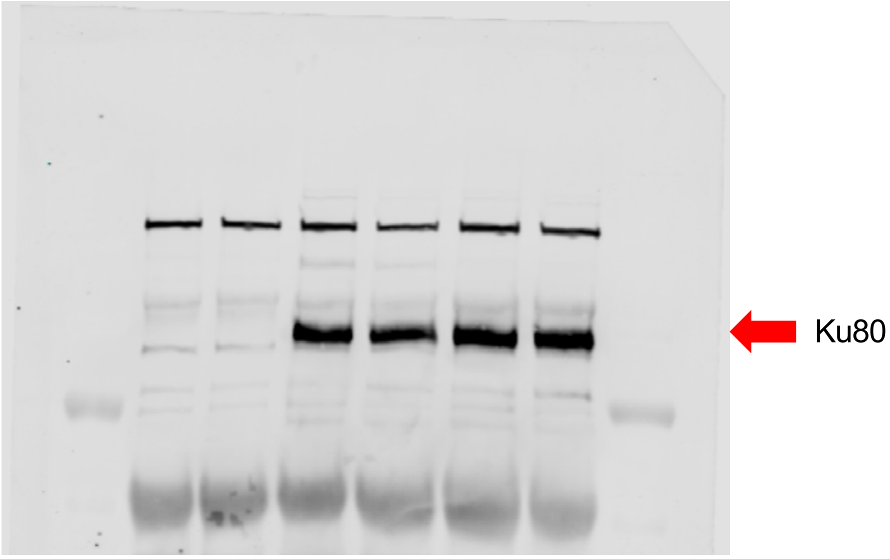


**Supplementary Figure S13.** Uncropped western blots shown in **Figure 5C.**

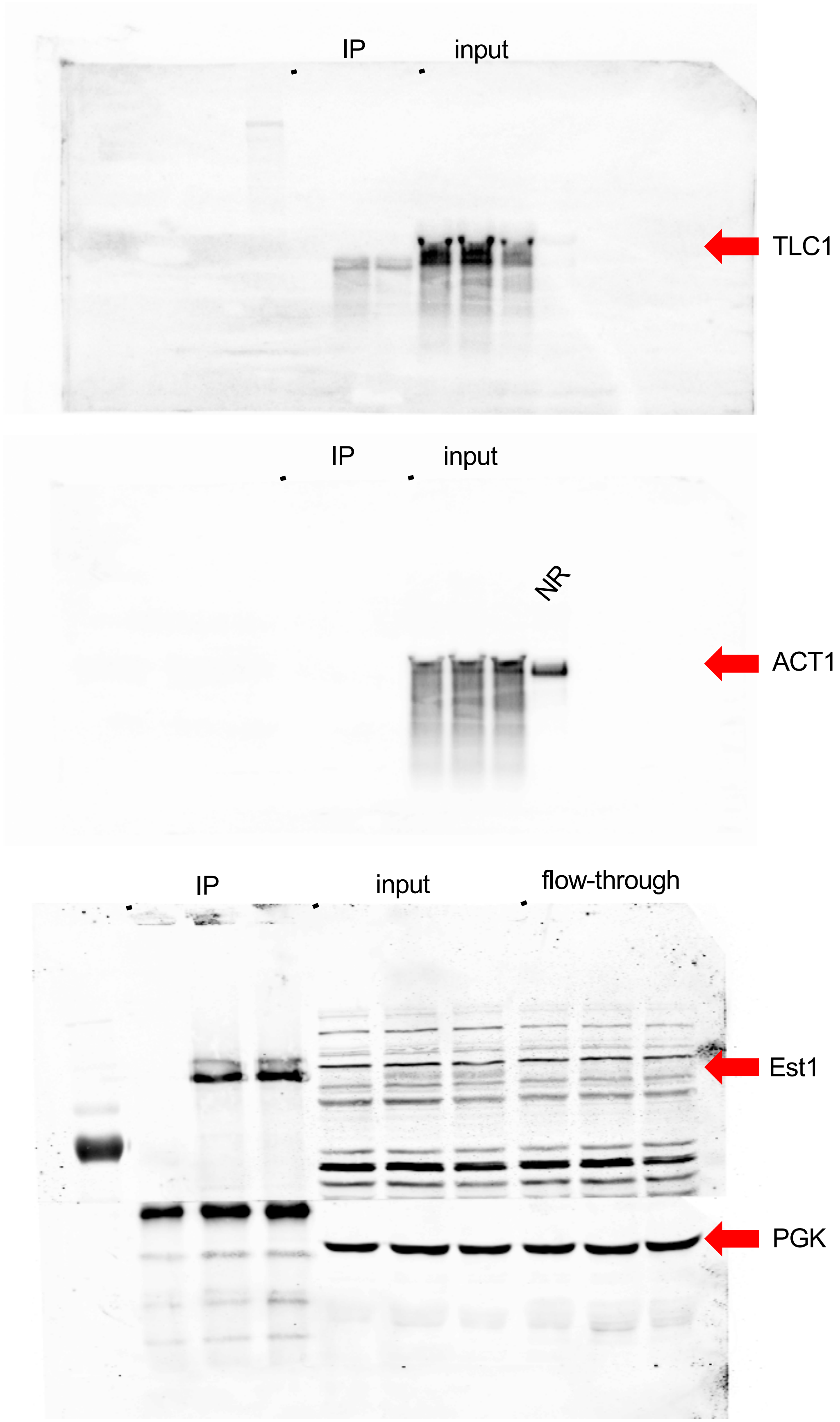


**Supplementary Figure S14.** Top: Uncropped northern blots shown in **Figure 5E**; NR indicates not relevant
Bottom: Uncropped western blots shown in **Figure 5E**

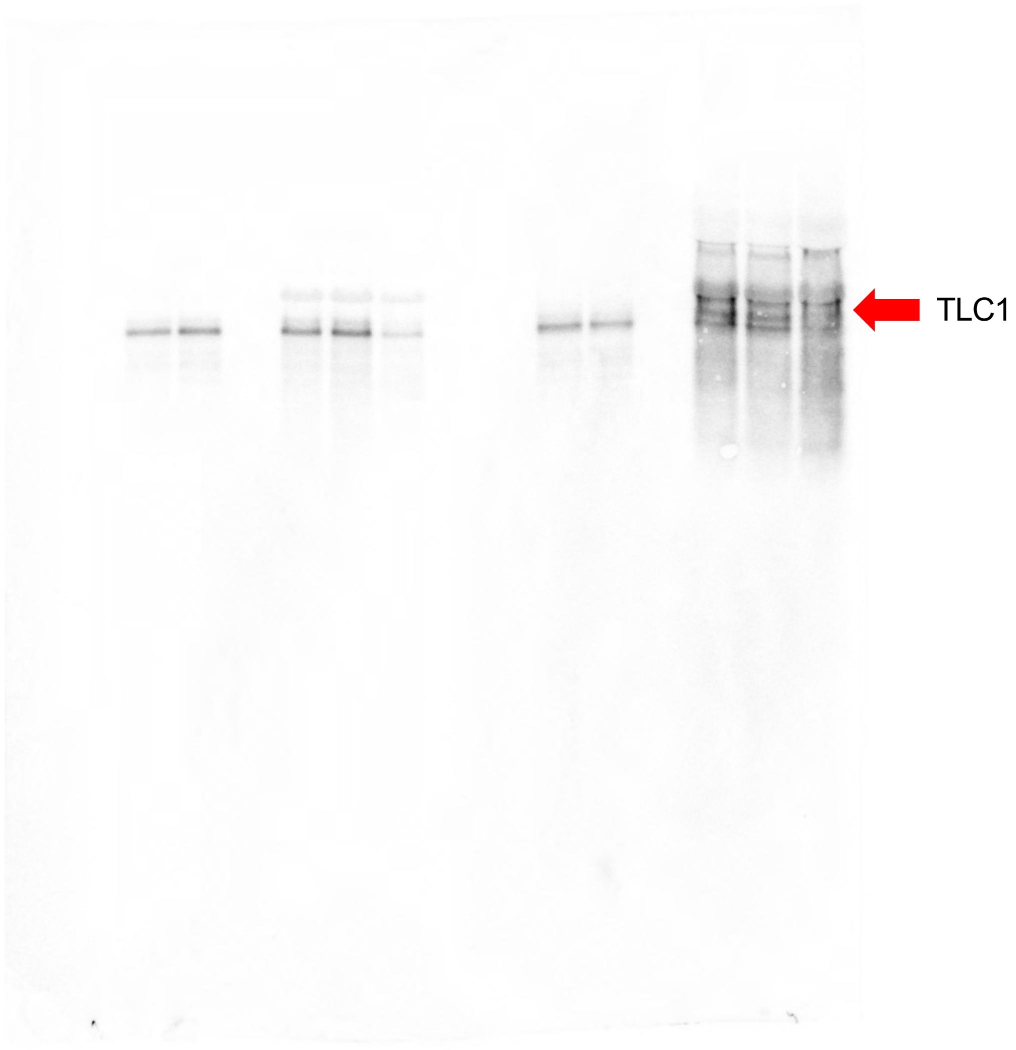


**Supplementary Figure S15A.** Uncropped northern blot shown in **Supplementary Figure S3,** which is representative of data quantified for **Figure 5H.**

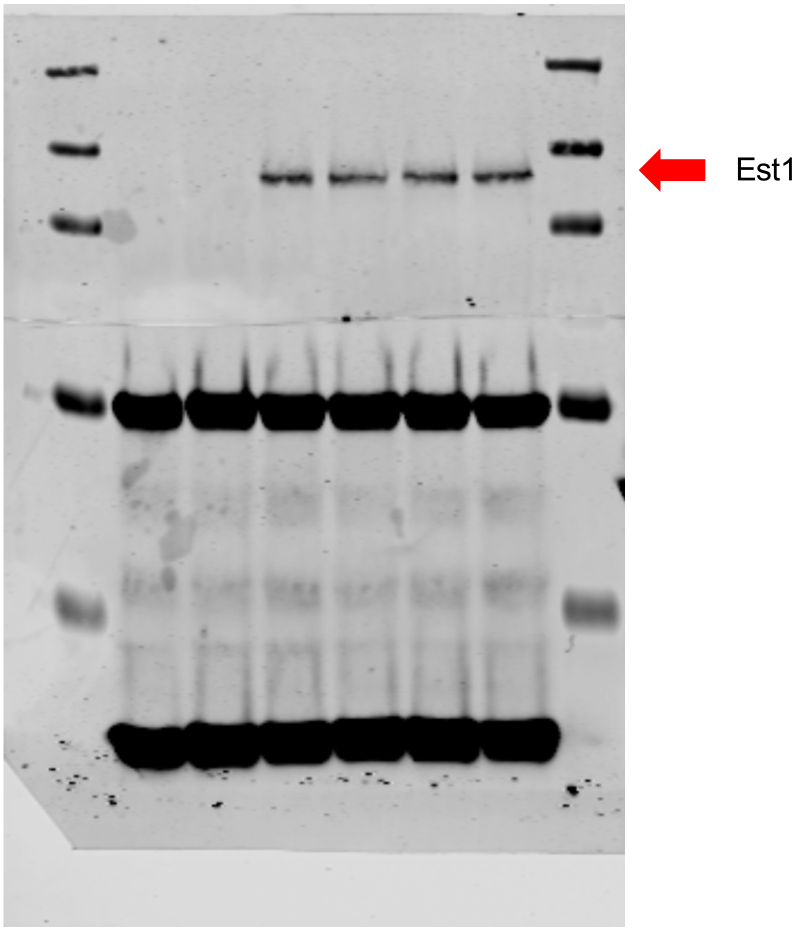


**Supplementary Figure S15B.** Uncropped western blot shown in **Supplementary Figure S3,** which is representative of data quantified for **Figure 5H.**

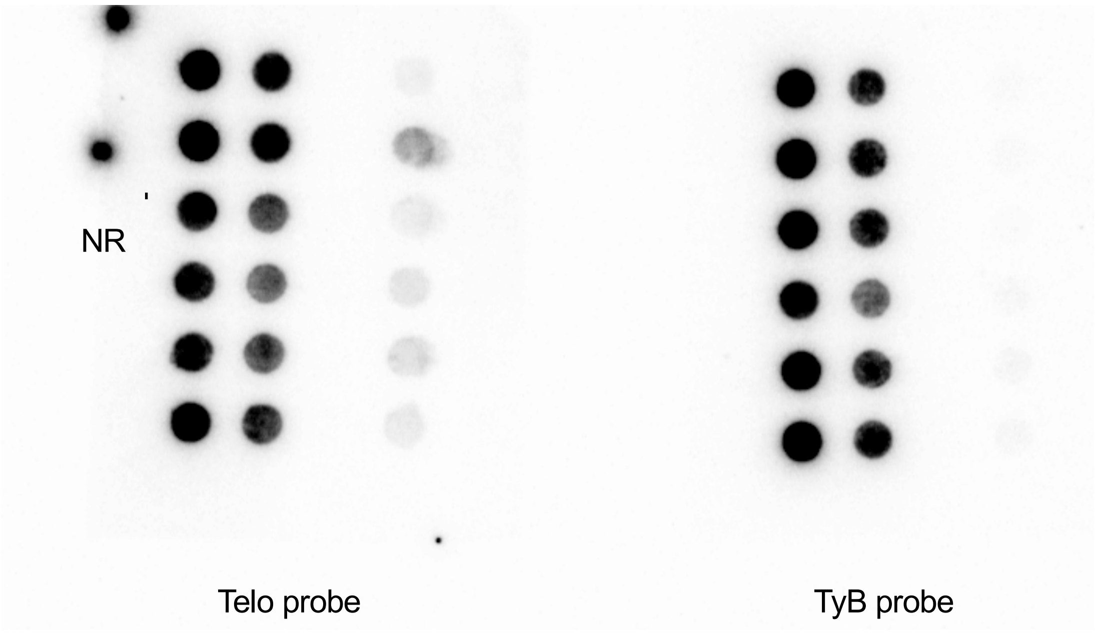


**Supplementary Figure S16.** Uncropped dot blots shown in **Figure 6A**; NR indicates not relevant.

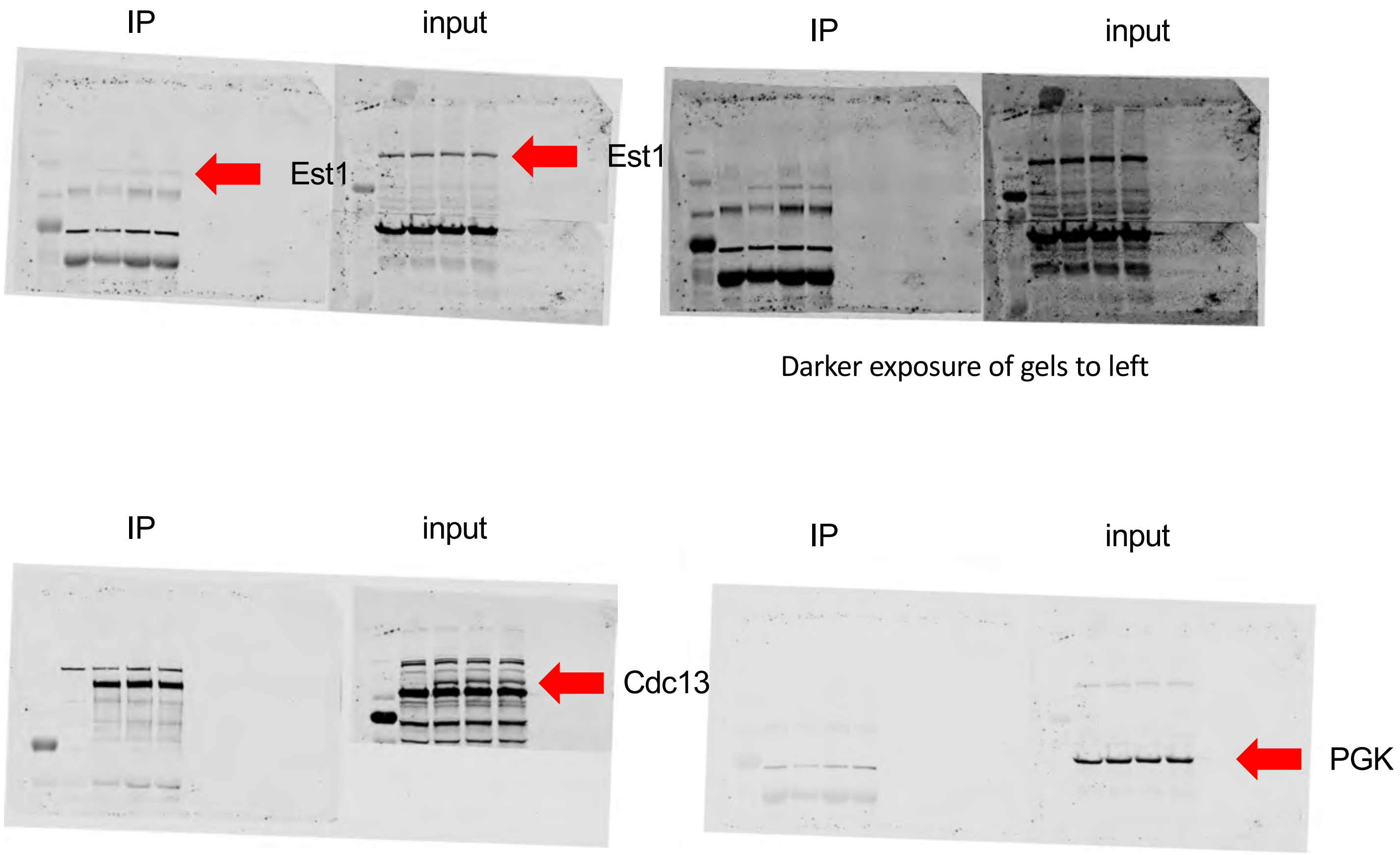


**Supplementary Figure S17A.** Uncropped top set of western blots shown in **Figure 7A**; NR indicates not relevant

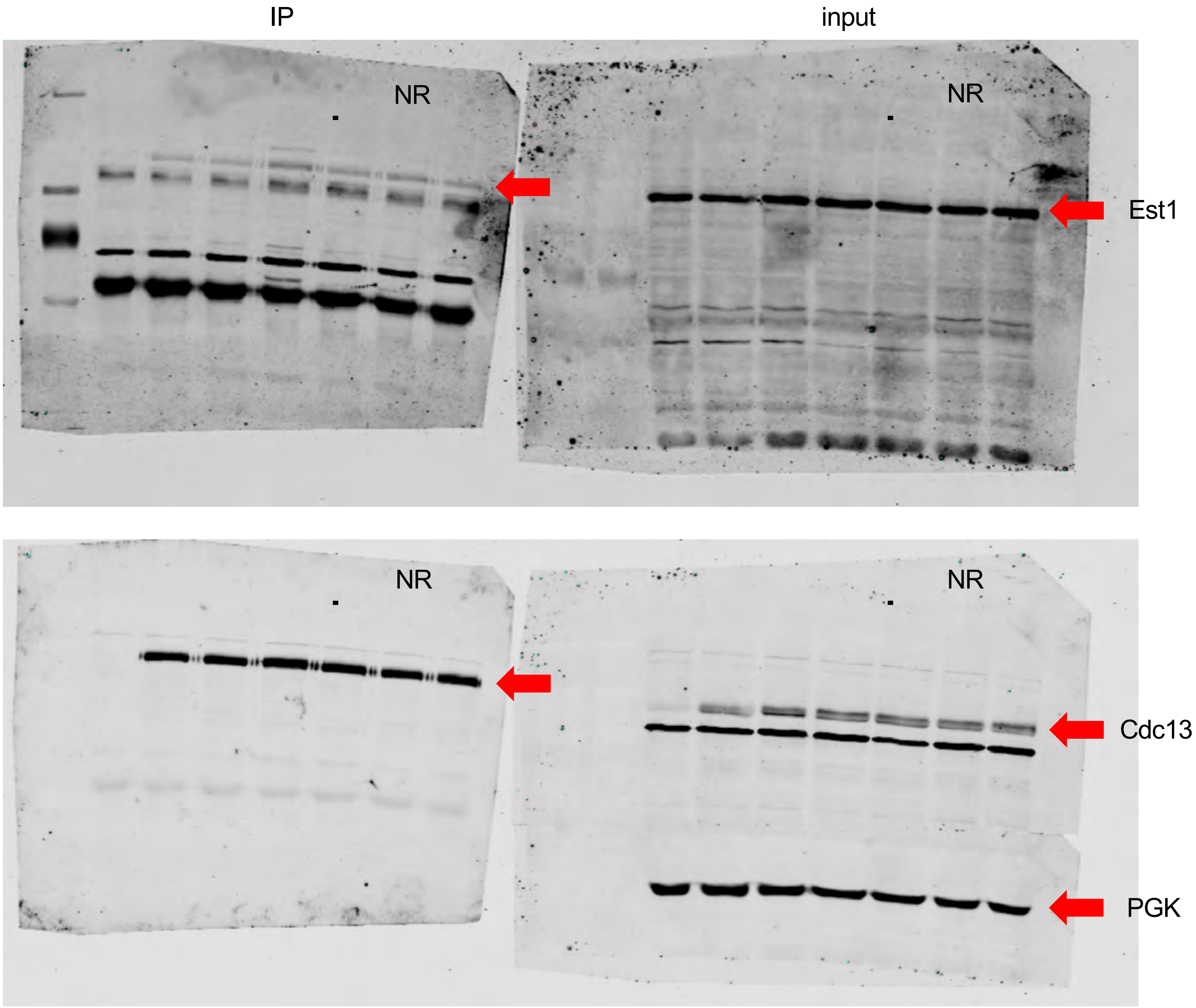


**Supplementary Figure S17B.** Uncropped bottom set of western blots shown in **Figure 7A**; NR indicates not relevant

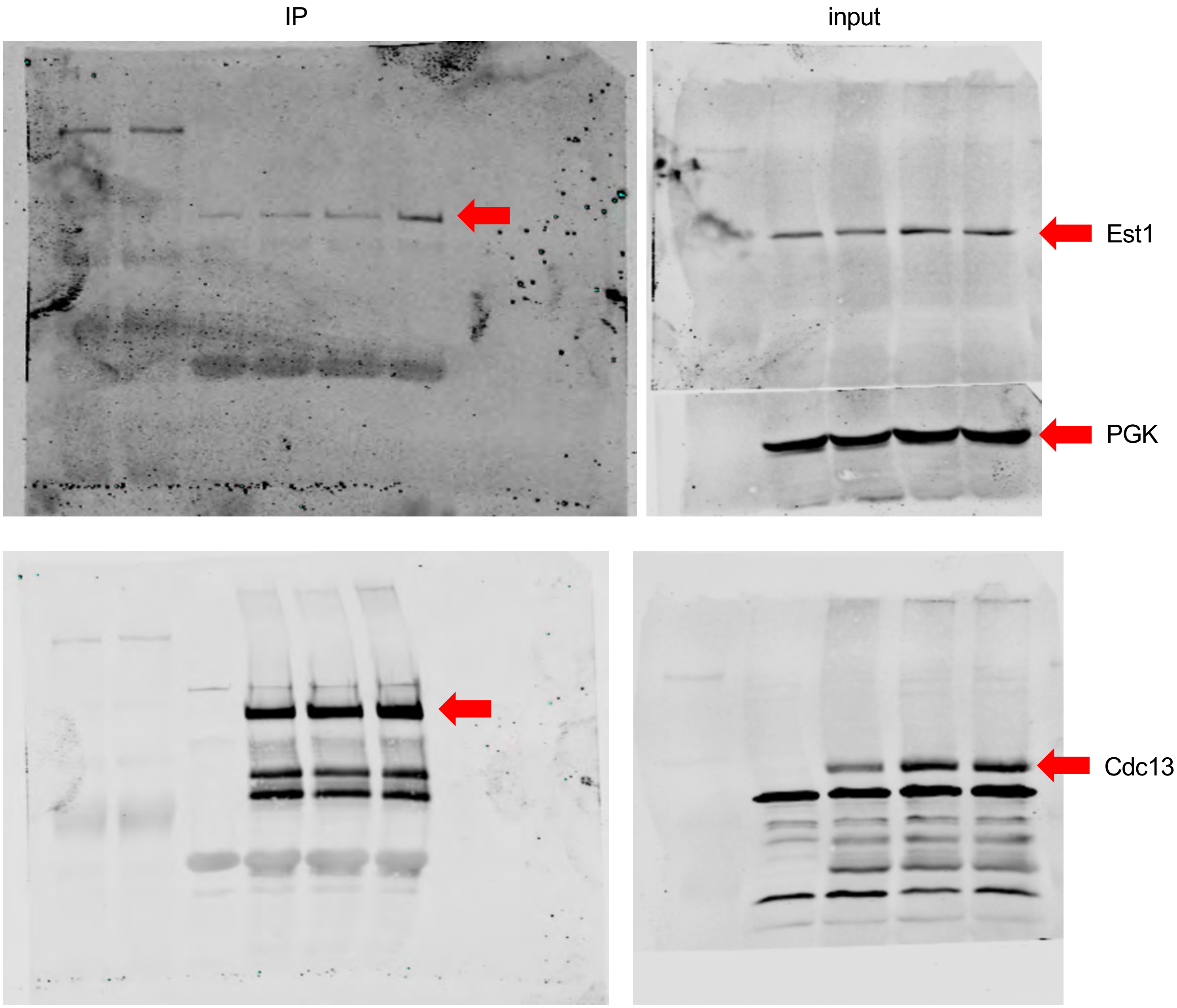


**Supplementary Figure S18**. Uncropped western blots shown in **Figure S4.**
